# Global gaps and priorities for shark and ray conservation: Integrating threat, function, and evolutionary distinctiveness

**DOI:** 10.1101/2025.11.28.691085

**Authors:** Théophile L. Mouton, Daniele Silvestro, Kristína Kocáková, Dominik Spitznagel, Fabien Leprieur, Catalina Pimiento

## Abstract

Elasmobranchs (sharks, rays, and skates) face unprecedented extinction risk, with over one-third of species threatened primarily by overfishing. While marine protected areas (MPAs) are essential tools to help safeguard marine ecosystems and the species that inhabit them, we lack a comprehensive understanding of how well elasmobranchs are protected globally. Here, we (1) evaluate the current level of protection for elasmobranchs, (2) employ CAPTAIN reinforcement learning algorithm to identify areas that optimise conservation based on species’ level of threat and their functional and evolutionary distinctiveness, and (3) examine conservation conflicts and opportunities by identifying areas where conservation priorities overlap with high and low fishing pressures. Our analysis revealed severe protection deficits, with elasmobranchs having on average only 3% of their range covered by no-take MPAs and 64% of species under-represented relative to random expectations. Areas that optimise conservation priorities consistently converged in Australian, Southeast Asia, the Indian Ocean, Atlantic Africa, European Atlantic coasts, the Caribbean, and South American Atlantic regions. However, areas of highest conservation priority often coincided with intensive fishing, particularly in East Asian seas, highlighting conservation conflicts requiring strategic management. Our results provide actionable insights for policymakers to optimise conservation strategies and maximise biodiversity retention.

**Teaser:** Reinforcement learning identifies priority areas to expand ocean protection for unique and irreplaceable elasmobranchs.

## 1. Introduction

Marine biodiversity is increasingly threatened by human activities, with overfishing representing the most pervasive pressure, spanning nearly the entire ocean surface and depth (Kroodsma et al. 2018; Welch et al. 2022; Jacquemont et al. 2024). This has pushed many vulnerable species to the brink of extinction (Dulvy et al. 2021; Yan et al. 2021). A striking example is elasmobranchs, which play critical ecological roles as predators and mesoconsumers, regulating ecosystems both directly through trophic interactions and indirectly by influencing prey behaviour (Dedman et al. 2024). Declines in elasmobranchs populations can profoundly disrupt ecosystem functioning and the services these ecosystems provide (Heithaus et al. 2008; Baum and Worm 2009).

According to the most recent global IUCN Red List of Threatened Species™ assessment (https://www.iucnredlist.org/), over one-third of elasmobranchs species are at risk of extinction, with three species possibly already extinct, marking what may be the first global marine fish extinctions caused by overfishing (Dulvy et al. 2021). This serial depletion began in rivers, estuaries, and coastal waters before expanding across the open ocean and into the deep sea (Dulvy et al. 2024). Consequently, there has been a significant loss of functionally important species, with estimates suggesting that the extinction of threatened species could lead to a 22% reduction in functional diversity (Dulvy et al. 2024). National extinction risk is particularly acute in countries with intense fishing pressure and substantial harmful fisheries subsidies, whereas nations with lower extinction risk tend to have stronger governance, larger economies, and greater investment in beneficial fisheries subsidies (Dulvy et al. 2024).

Protected areas are widely regarded as a cornerstone for conserving threatened ecosystems, with global targets set by the Kunming-Montreal Global Biodiversity Framework aiming to protect 30% of land and oceans by 2030, including 10% under strict protection (CBD 2022). In response, the global designation of protected areas has accelerated, but the rate of expansion has varied significantly across nations and regions over recent decades (Farhadinia et al. 2022; Mouton et al. 2025). Despite the adoption of this target by 196 parties, there remains no consensus on prioritising sites for protection or identifying which ecosystems or habitat types are most in need of safeguarding (Aminian-Biquet et al. 2025; Watson et al. 2023). In marine environments, marine protected areas (MPAs) have been increasingly recognised not only for enhancing fisheries yields and spillover profits (Di Lorenzo et al. 2020; Medoff et al. 2022) but also for providing critical benefits such as climate change mitigation and resilience (Jacquemont et al. 2022; Roberts et al. 2017).

In some localised areas of the world, MPAs have demonstrated benefits for shark populations, including endemic and threatened species off South Africa (Albano et al. 2021), Caribbean Reef Sharks in Belize (Bond et al. 2017), coastal sharks in the Eastern Pacific (Klimley et al. 2022), and Grey Reef Sharks in Western Australia (Speed et al. 2018). However, the effectiveness of MPAs in conserving wide-ranging and migratory species remains limited due to the relatively small area they cover (Dwyer et al. 2020). Furthermore, the extent to which the geographic ranges of the most imperilled shark species are covered by MPAs remains poorly understood, and there is still no consensus on which areas should be prioritised to maximise conservation outcomes.

Beyond threat status alone, conservation planning increasingly recognises the importance of preserving multiple dimensions of conservation value. Functional diversity captures the variety of ecological roles species perform, with functionally unique species providing irreplaceable ecosystem services that cannot be compensated by other taxa (Mouillot et al. 2013; Petchey and Gaston 2006). Evolutionary distinctiveness measures the uniqueness of a species’ evolutionary history, with phylogenetically distinct lineages harbouring irreplaceable genetic heritage and adaptive potential (Isaac et al. 2007). Approaches like FUSE (Functionally Unique, Specialised, and Endangered) and EDGE (Evolutionary Distinct and Globally Endangered) integrate these dimensions with extinction risk to identify species whose loss would be most consequential for maintaining both ecological functions and evolutionary diversity (Pimiento et al. 2020; Isaac et al. 2007). Much of the functional and phylogenetic diversity of elasmobranch assemblages falls outside existing MPA networks (Pimiento et al. 2023), yet no species-level assessment has integrated threat-based conservation priorities with functional and evolutionary distinctiveness dimensions for elasmobranchs at the global scale. This gap limits our ability to develop comprehensive conservation strategies that protect not only the most threatened species, but also those most critical for maintaining ecosystem functioning and evolutionary potential (Thuiller et al. 2015).

With finite conservation resources and competing demands for marine space, systematic conservation prioritisation is essential for identifying areas that can maximise biodiversity protection while balancing ecological, economic, and social constraints. Several tools like Marxan (Ball et al. 2009), Zonation (Moilanen et al. 2009), and prioritizeR (Hanson et al. 2024) have been instrumental in guiding conservation decisions. Marxan is widely used for cost-effective reserve design, optimising networks of protected areas to meet biodiversity targets under budgetary constraints (e.g. Dwyer et al. 2019). Zonation, on the other hand, ranks entire landscapes based on their contribution to biodiversity retention, making it ideal for continuous prioritisation across large areas (e.g. Pollock et al. 2017). Similarly, prioritizeR offers flexibility in incorporating advanced statistical modelling and optimisation frameworks, particularly useful for researchers requiring customisation and reproducibility. Another such tool is the CAPTAIN algorithm (Conservation Area Prioritisation Through Artificial Intelligence Networks; Silvestro et al. (2022)) which uses reinforcement learning to train neural networks that identify conservation priority areas by analysing species distribution patterns to maximise user-defined conservation objectives (Silvestro et al. 2025).

Here, we identify global priority areas to optimise the conservation of different facets of elasmobranch biodiversity. To do so, we first assess current protection gaps by quantifying the spatial overlap between species ranges and existing marine protected area (MPA) networks. Then, we employ the CAPTAIN algorithm under a predefined budget of 10% no-take MPAs, to identify priority areas to safeguard (1) endangered species, (2) Functionally Unique Specialised and Endangered (FUSE) species and (3) Evolutionary Distinct and Globally Endangered (EDGE2) species. Finally, we identify both regions where high conservation priorities coincide with intensive fishing, and areas offering proactive conservation opportunities. By combining gap analysis, systematic prioritisation, and conflict assessment, we aim to provide actionable insights for policymakers to optimise conservation strategies and maximise biodiversity retention.

## 2. Methods

### 2.1 Elasmobranch distributions

We used the global range maps for elasmobranchs (sharks, rays, and skates) from Pimiento et al. (2023) to assess species distributions. These maps, based on species-level data from the IUCN Red List of Threatened Species (www.iucnredlist.org; last accessed October 2021), were mapped onto a global 0.5° resolution grid to create a presence/absence matrix of 1,075 species across 131,327 grid cells.

### 2.2 Dimensions of conservation value

To optimise conservation prioritisation for elasmobranchs, we used three complementary biodiversity metrics: (1) IUCN Red List threat status (hereafter referred to as IUCN status), (2) Functionally Unique Specialised and Endangered (FUSE) scores (Pimiento et al. 2020), and (3) Evolutionary Distinct and Globally Endangered (EDGE2) scores (Stein et al. 2018; Isaac et al. 2007; Gumbs et al. 2023). Values for all three metrics were extracted from Pimiento et al. (2023).

IUCN threat status was treated as an ordinal variable with five categories (1 = Least Concern, 2 = Near Threatened, 3 = Vulnerable, 4 = Endangered, 5 = Critically Endangered). FUSE is calculated based on seven ecological and morphological traits: maximum body size (total length or disk width), habitat (coastal, oceanic, or both), salinity tolerance (marine, brackish, or freshwater), vertical position (benthic, benthopelagic, or pelagic), prey preference (vertebrates, fish, invertebrates, plankton, or combinations), feeding mechanism (macropredators or filter feeders), and thermoregulation (ectothermic or mesothermic). A trait dissimilarity matrix is constructed using a modified Gower’s distance to accommodate different variable types while ensuring equal trait weighting (Pavoine et al. 2009). Principal Coordinate Analysis (PCoA) is performed on this matrix to construct a multidimensional functional space, from which Functional Specialisation (FSp; measuring niche breadth) and Functional Uniqueness (FUn; measuring trait dissimilarity from other species) are calculated. The FUSE metric integrates these two functional components with extinction risk (IUCN status) into a single composite score.

EDGE2 incorporates evolutionary distinctiveness (ED2) calculated from time-calibrated phylogenies, weighted by the extinction probabilities of each species and its relatives, equivalent to the Heightened Evolutionary Distinctiveness (HED) metric.

After integrating these conservation metrics with spatial distribution data, our subsequent analyses focused on 1,000 species with complete datasets across all components (traits, spatial distributions, phylogeny, and conservation metrics), excluding exclusively freshwater species to maintain consistency with the marine and brackish focus of our conservation assessments.

### 2.3. Marine Protected Areas

We downloaded a global shapefile of marine protected area location from the World Database on Protected Areas (https://www.protectedplanet.net/en/thematic-areas/wdpa?tab=WDPA) in September 2024. We considered a single group of MPAs according to their level of protection: no-take MPAs (where any type of extractive activity is forbidden; i.e., those aligning with IUCN categories I, II, or III; following Day et al. (2019)).

### 2.4 Spatial gap analysis

To assess the coverage of Marine Protected Areas (MPAs) across shark and ray species ranges, we conducted a spatial gap analysis using R (version 4.4.2; R Core Team 2024). Species occurrence data were mapped onto a global grid, and we calculated the overlap between species ranges and no-take MPAs, using binary raster layers. For each species, we quantified the percentage of its range that falls within protected areas by calculating the ratio of grid cells that overlap with MPAs to the total number of cells in the species’ range.

We assessed the relationship between IUCN threat status and current MPA coverage using non-parametric approaches due to violations of parametric assumptions. Prior to analysis, we tested key assumptions for linear models. Levene’s test, computed using the *car* package (Fox & Weisberg, 2018) revealed significant heterogeneity of variance across threat categories (F = 7.47, p < 0.001), and Shapiro-Wilk tests indicated strong departures from normality (W = 0.55, p < 0.001). These violations were driven by the highly right-skewed distribution of coverage values (skewness: 2.1-5.0 across categories, calculated using the *moments* package; Komsta & Novomestky, 2015) and high proportion of species with zero coverage (29-35% per category). Given these violated assumptions, we used the Jonckheere-Terpstra test for ordered alternatives, using the *DescTools* package (Signorell et al., 2024) to assess the presence of a monotonic declining trend across threat categories. This non-parametric test is specifically designed for ordered categorical predictors and is robust to skewed distributions and heterogeneous variances (Jonckheere 1954; Terpstra 1952). We corroborated this finding using Spearman’s rank correlation and the Kolmogorov-Smirnov two-sample test comparing extreme categories (LC vs CR), both computed using the *stats* package (R Core Team 2024). To examine heterogeneity in this relationship across the distribution of coverage values, we also conducted quantile regression for the 30th through 90th percentiles using the *quantreg* package (Koenker, 2024). Standard errors were estimated using bootstrap resampling with 500 replications (Koenker, 2005). Quantile regression is appropriate for examining conditional quantiles when relationships may vary across the response distribution, particularly with highly skewed data at distributional boundaries (Cade & Noon, 2003).

To evaluate whether the observed MPA coverage differed from what would be expected by chance, we developed a spatially explicit null model that accounted for both marine areas and exclusive economic zones (EEZs). The null model randomly placed MPAs within each country’s EEZ while maintaining the same total area of protected space per country as observed in the actual MPA network. We ran 100 iterations of this randomisation procedure, using parallel processing for computational efficiency. For each species, we calculated the mean and standard deviation of MPA coverage across all iterations, separately for both all MPAs and no-take MPAs. We then compared observed coverage against null model expectations by calculating standardized effect sizes (z-scores; SES) for each species, defining significant over- or under-representation using a threshold of ±1.96 standard deviations from the null expectation (corresponding to a 95% confidence interval). Species were classified as over-represented if their observed MPA coverage was significantly higher than expected by chance, and under-represented if significantly lower. To visualise spatial patterns in protection, we mapped the mean standardized effect sizes across all species present in each grid cell, using a diverging colour scale centred at zero.

### 2.5 Conservation prioritisation

To optimise spatial conservation prioritisation, we employed the CAPTAIN framework (Silvestro et al. 2022), using the latest development version 2.3 of the software in Python 3.12.7 (Python Software Foundation, https://www.python.org). CAPTAIN maps the spatial distribution of features such as species distributions and their extinction risks onto a selection of conservation priorities using a neural network. The network is optimised within a reinforcement learning framework aiming to maximize a user-defined reward function. The selection of areas for protection can be further constrained by budgetary constraints or by a defined target fraction of protected area.

Here we optimised three models with reward functions based on IUCN status, FUSE and EDGE2, respectively. The IUCN status data were treated as five classes ranging from 1 for Least Concern species to 5 for Critically endangered species. Similarly, we discretised the FUSE and EDGE2 metrics into 5 classes, where class 1 was associated with the lowest scores and class 5 with the highest scores. To each class we assigned weights used in the reward function such that highest priority is given to species with the highest weight (Silvestro et al. 2025):

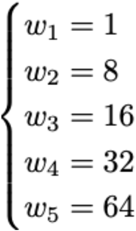

The reward functions were defined as

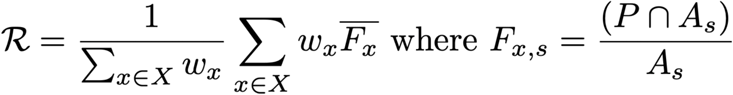

where X = {1, …, 5} is the score associated with each species (based on IUCN status, FUSE or EDGE2), F_x, s_ is the proportion of geographic range of species *s* (A_s_) included in protected areas (P). The value

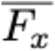

is the proportion of protected range averaged across all species with score *x*. The reward is therefore a weighted average of the fraction of protected range achieved across all species.

We trained the model using the elasmobranch data with unlimited budget and a target of 10% of the cells in our 0.5° resolution presence/absence matrix. This budget was used because the Global Biodiversity Framework’s Target 3 calls for the conservation of at least 30% of marine areas by 2030, with at least 10% of these areas under strict protection. The three models with different reward functions were trained for 500 epochs using a batch size of 4. The trained models were then used to perform 50 predictions based on the network weights of the last 50 epochs. We used the frequency at which each cell was selected to be protected as a measure of priority in follow up analyses. The performance of CAPTAIN was assessed in terms of species preserved and areas protected.

To evaluate whether CAPTAIN’s spatial priorities effectively targeted species according to each dimension’s underlying metric, we examined the relationship between species-level protection outcomes and dimension of conservation value scores. This analysis served as a validation step to confirm the prioritisation algorithm functioned as intended before interpreting spatial patterns. For each species, we calculated the percentage of its geographic range that fell within priority areas at the 10% budget level for each of the three dimensions (IUCN status, FUSE, and EDGE2). We then analysed the relationship between these protection percentages and their respective scores: IUCN status (ordinal categories 1-5: LC, NT, VU, EN, CR), FUSE scores (discretised categories 1-5), and EDGE2 scores (discretised categories 1-5).

We examined these relationships using beta regression models (Ferrari & Cribari-Neto, 2004), which are appropriate for bounded percentage data. Protection percentages were transformed to the (0,1) interval using the Smithson & Verkuilen (2006) boundary correction: y’ = (y × (n - 1) + 0.5) / n, where y is the proportion and n is the sample size.

Prior to model fitting, we assessed key assumptions through comprehensive diagnostic tests. Levene’s tests showed homogeneous variance for the IUCN model (F = 2.19, p = 0.07) but heterogeneity for FUSE and EDGE2 models (F = 5.42-8.72, p < 0.001). Shapiro-Wilk tests on deviance residuals indicated departures from normality (W = 0.93-0.95, all p < 0.001), though these violations were substantially milder than those observed in current coverage analysis (W = 0.87). Critically, CAPTAIN-generated protection distributions showed near-symmetric distributions (skewness: −0.23 to 0.29) with minimal zero inflation (6.5-11.7% of species) and few influential observations (1.6-2.3% exceeding Cook’s distance threshold), contrasting sharply with the highly skewed current coverage data (skewness: 2.1-5.0, 29-35% zeros, 4.3% influential observations). Given these substantially improved distributional properties and beta regression’s specific design for bounded data, we deemed beta regression appropriate for quantifying CAPTAIN’s effectiveness.

We fit separate beta regression models for each dimension, with species-level protection as the response variable and the respective dimension score as the predictor. We report coefficients on the logit scale, standard errors, z-values, p-values, pseudo-R² values, and average marginal effects (AMEs) calculated using the *margins* package (Leeper et al., 2021). AMEs quantify the average change in the percentage of species’ range protected for each unit increase in dimension score. Beta regression analyses were conducted using the *betareg* package (Zeileis et al., 2016).

### 2.7 Spatial distribution analysis

To quantify the spatial distribution patterns of high-priority areas across the three prioritisation dimensions, we calculated mean nearest neighbour distances (MNN) for cells with priority values >0.9. MNN measures the average distance between each high-priority cell and its closest neighbouring high-priority cell, providing a metric of spatial clustering versus dispersion. For each dimension (IUCN, FUSE, and EDGE2), we extracted the centroids of high-priority cells and computed pairwise nearest neighbour distances using the *spatstat* package (Baddeley et al. 2014). Lower MNN values indicate more spatially clustered priority areas, while higher values indicate more geographically dispersed patterns. We report both mean and standard deviation of nearest neighbour distances to characterise the central tendency and variability in spatial clustering. The analysis was conducted in the McBryde-Thomas Flat-Polar Sinusoidal projection (+proj=mbt_s), with distances measured in kilometres.

### 2.8 Spatial congruence analyses

We developed a statistical framework to assess spatial congruence between the three prioritisation outcomes that can be applied across any priority threshold value, following Albouy et al. (2015). For our analysis, we selected a threshold of 0.9 as it represents an optimal balance between identifying genuinely high-priority areas while maintaining sufficient spatial coverage for meaningful conservation planning. The analysis quantifies the percentage of overlap as the ratio between the observed overlap (number of cells classified as high-priority in both indices) and the minimum number of high-priority cells between the two indices, providing a conservative estimate of spatial congruence. We assessed the statistical significance of the observed overlap using a randomisation test with 999 permutations. For each permutation, we randomly reshuffled the spatial distribution of one dimension’s high-priority areas while maintaining the other fixed and calculated the resulting overlap. The p-value was computed as the proportion of random overlaps that were more extreme (larger if observed > expected, smaller if observed < expected) than the observed overlap.

### 2.9. Fishing pressure and conservation priorities

To identify potential conflicts between fishing activities and conservation priorities, as well as opportunities for proactive protection, we estimated global fishing effort using two complementary data sources. First, we utilised the latest version of fleet daily fishing activity data (Kroodsma et al. 2018) from Global Fishing Watch (GFW) with the finest available resolution (0.1-degree). GFW compiles data from publicly accessible automatic identification systems (AIS) and government-operated vessel monitoring systems (VMS). The dataset is based on fishing detections from over 114,000 unique AIS-equipped fishing vessels, with approximately 70,000 active each year. Although only 2% of all fishing vessels are equipped with AIS (primarily large industrial vessels), these vessels account for 50% of fishing activity within exclusive economic zones (Kroodsma et al. 2018). We used data from the four most recent years available on the GFW data download platform as of September 2024: 2017 to 2020. Although this dataset is unique and represents the best available estimate of global fishing effort, in some regions of the world, vessels do not use AIS nor VMS. This is particularly the case in the Caribbean Sea, the southern Mediterranean Sea, the Northern Arabian Sea including the Red Sea and Gulf of Oman, and the seas of the Coral Triangle.

For this reason, we used the latest industrial vessel detections from Sentinel-1 Synthetic Aperture Radar (SAR; Paolo et al. (2024)). Sentinel-1 SAR is uniquely suited for detecting vessels at sea, as it operates independently of light conditions and cloud cover, allowing for consistent monitoring in all weather conditions, day or night. By utilising SAR imagery, Paolo et al. (2024) detected vessels that are not required to broadcast their location via AIS or VMS, especially in regions where “dark vessels” (those that turn off their transponders or do not have them) are prevalent. The Sentinel-1 SAR data can detect vessels larger than 15 meters, with a detection rate exceeding 70% for vessels around 25 meters in length and over 90% for vessels 50 meters and larger. It is concentrated in inshore waters (< 200 meters depth), mostly due to satellite coverage. We used monthly data from January 2017 to December 2020 to match fleet daily fishing activity data and summed the number of fishing vessel detections in each 0.1 degree cell. Fishing was defined as vessel detections that had a fishing probability index of 0.9 or more (Paolo et al. 2024). The fishing probability index is derived from combining AIS data, Sentinel-1 SAR imagery, and deep-learning models to identify fishing vessels that were not broadcasting AIS signals.

From this integration, we had 87.6% of grid cells with only AIS data, 9.1% with both AIS and SAR and 3.2% with only SAR data. We built a random forest model from > 160,000 associations between AIS and SAR detections, with the *randomForest* package in R (Breiman et al. 2018), to predict fishing hours in areas with only SAR detections. Fishing hours were log + 1 transformed and the distance to ports, distance to shore, bathymetry (all three layers obtained from Global Fishing Watch) and geographical coordinates (latitude and longitude) used as additional predictors in the model. This model performed well, with 82.4% of variation explained and a Mean Absolute Error of 186.5 (Median Absolute Error = 10.1; Supplementary methods). These predictions allowed us to obtain estimates of fishing hours in parts of the Caribbean Sea, the Southern Mediterranean Sea and coastal areas of the Coral Triangle. However, there were still major areas without any estimate of fishing effort: 56.5% of all 0.1-degree resolution ocean grid cells, composed mostly of polar waters but also major areas of the Caribbean Sea, the Northern Arabian Sea, the Bay of Bengal and the Coral Triangle.

This gap in estimates of fishing effort was a major limitation to the use of CAPTAIN, as preliminary runs of the model consistently prioritised cells without estimates of fishing effort. Consequently, we excluded the fishing effort layer from our CAPTAIN analyses to prevent systematic bias towards data-deficient areas in conservation prioritisation.

To assess conservation conflicts and opportunities, we examined the relationship between conservation priorities and fishing pressure using two complementary approaches. First, we created bivariate maps overlaying cell-level priority values from each dimension (IUCN, FUSE, and EDGE2) with cumulative commercial fishing effort from 2017-2020, allowing visual identification of spatial overlaps between high conservation priorities and intensive fishing pressure. Second, we aggregated these patterns at the ecoregional level using the Marine Ecoregions of the World (MEOW) classification system. For each marine ecoregion, we calculated mean conservation priority scores from our three prioritisation indices and mean fishing effort (total fishing hours) across all grid cells within ecoregion boundaries. This ecoregional analysis allowed us to quantify relationships between conservation priorities and fishing pressure, identifying regions where high conservation priorities coincide with intensive fishing pressure (indicating conservation conflicts requiring immediate intervention) versus areas where high conservation values occur under low fishing pressure (representing proactive conservation opportunities). We visualised ecoregion-level relationships through scatterplots of mean fishing effort versus mean conservation priority, with ecoregions coloured by biogeographic realm to highlight regional patterns.

## 3. Results and Discussion

### 3.1 Gap analysis of no-take marine protected areas

To evaluate how well current no-take MPAs cover the geographic distribution of elasmobranchs across threat categories, we calculated species-level range overlaps with no-take MPAs. We found overlaps ranging from 0.00% to 50.00%, with an average of 2.57% (sd = 5.39%). Our results align with regional assessments showing that Important Shark and Ray Areas overlap with no-take MPAs by only 7.3% in the Central and South American Pacific (Mouton et al. 2025). Further assessments at regional and national scales are needed to identify specific contexts where elasmobranch protection is adequate versus critically deficient, and to understand the drivers of these disparities.

Mean MPA coverage showed a significant declining trend across IUCN threat categories (Jonckheere-Terpstra test for ordered alternatives: J-T = 147,97, p < 0.01), with Least Concern species having 3.63% mean coverage, followed by Near Threatened (2.03%), Vulnerable (2.07%), Endangered (1.58%), and Critically Endangered species having the lowest coverage (1.41%; Fig. 1A). This pattern was corroborated by Spearman’s rank correlation (ρ = −0.08, p < 0.05) and pairwise comparison of extreme categories (Kolmogorov-Smirnov test, LC vs CR: D = 0.22, p < 0.01).

**Fig. 1.**
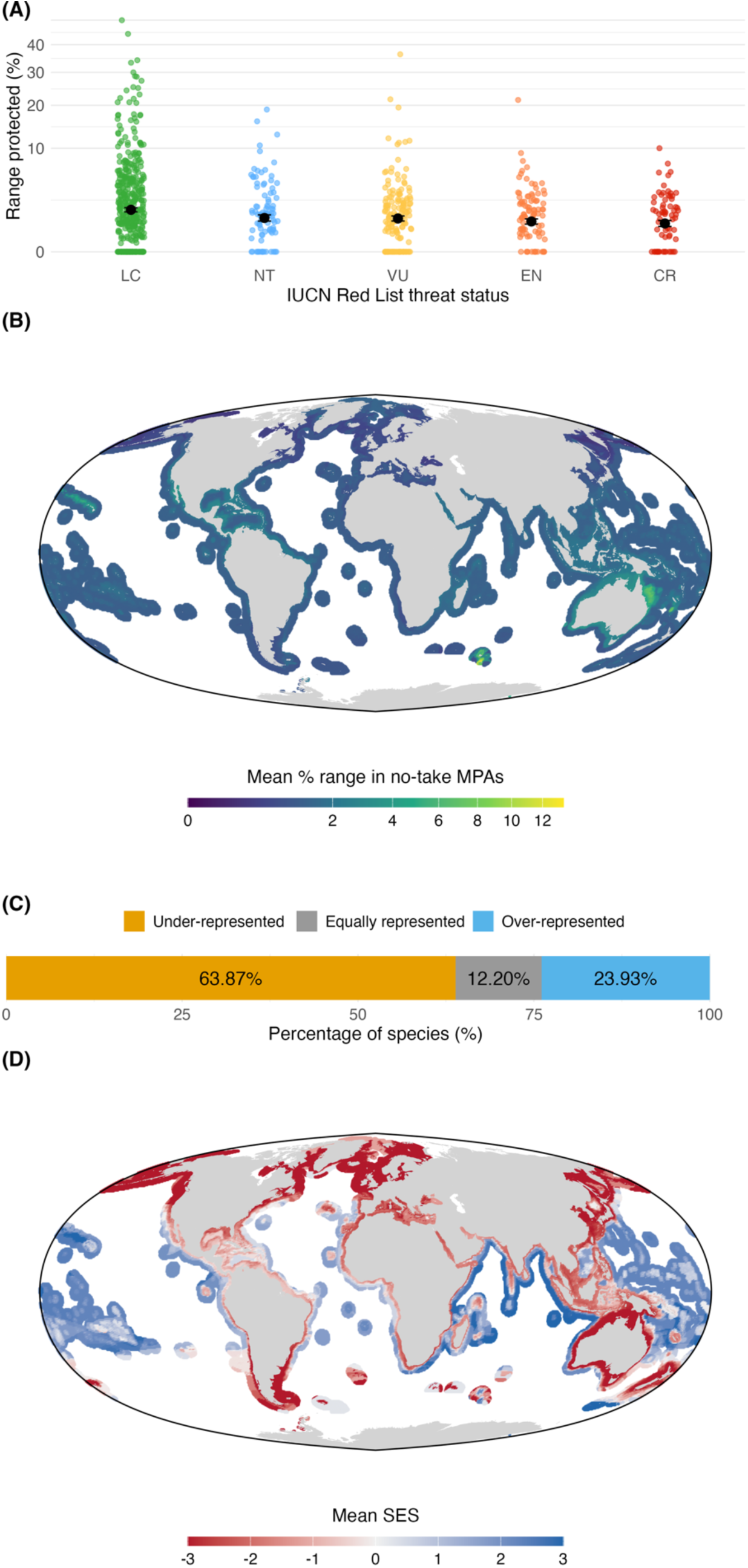
Marine protected area coverage and conservation status of global elasmobranch fauna. **(A)** Relationship between IUCN status and no-take MPA coverage for elasmobranch species. Scatter plot showing the percentage of species’ geographical range within no-take Marine Protected Areas (MPAs) across different IUCN threat categories. The y-axis is square root transformed to improve visibility of the distribution. Black dots represent mean values with error bars showing ±1 standard error. Coloured points represent individual species with IUCN status-specific colouring (LC=green, NT=light blue, VU=yellow, EN=orange, CR=red). **(B)** Global map of continental waters showing the mean percentage of elasmobranch species’ geographical ranges overlapped by no-take MPAs. Data are aggregated into a 0.5° x 0.5° grid, with colours indicating the mean percentage overlap in each grid cell. Mean percentage overlap is calculated only for cells containing species presence, with darker colours representing lower overlap. Land areas are shown in light grey, and grid cells are coloured on a square-root scale for better visualisation. **(C)** Representation of global elasmobranch fauna in no-take MPAs relative to random placement expectations. The horizontal stacked bar shows the percentage of elasmobranch species that are under-represented (orange, 63.91%), equally represented (grey, 12.19%), and over-represented (blue, 23.91%) in MPAs compared to a null model of MPA distribution. Under-represented species have less of their range overlapping with MPAs than expected by chance, equally represented species have range overlap consistent with random MPA placement, and over-represented species have greater range overlap with MPAs than expected by chance. **(D)** Spatial distribution of mean Standardized Effect Size (SES) per grid cell, showing the average deviation between observed and expected no-take MPA coverage across species ranges. Red colours indicate areas where species have, on average, lower MPA coverage than expected by chance (negative mean SES), while blue colours indicate higher coverage than expected (positive mean SES). Grey areas represent land masses. SES values were calculated for each species by comparing observed MPA coverage to null models, then averaged across all species present in each grid cell. The diverging colour scale is centred at zero, where observed coverage matches expectation.

Quantile regression clarified the nature of this relationship (Table S1; Fig. S1). While all slopes were negative from the 40th through 90th percentiles (slopes = −0.01 to −1.38) and effectively zero at the 30th percentile (slope = 0.00), both the magnitude of the decline and its statistical significance increased monotonically with quantile level (Table S1). The declining trend was only statistically significant for species in the upper quantiles of protection (70th-90th percentiles: slopes = −0.39 to −1.38, all p < 0.001), with no significant relationship detected for species with lower baseline protection (30th-60th percentiles: slopes = −0.01 to −0.14, p > 0.11). This indicates that while species are generally poorly protected regardless of threat status, among those that do receive protection, non-threatened species tend to have substantially higher coverage than threatened species, with this disparity becoming increasingly pronounced at higher protection levels.

To examine spatial patterns in protection coverage and identify priority gaps, we calculated, for each ocean grid cell, the mean percentage of species’ global ranges that overlap with no-take MPAs, averaging across all species present in that cell. The mean percentage range overlapped by no-take MPAs varied significantly across continental ocean grid cells, reaching a maximum of 13.44% and a minimum of 0.00% (mean = 1.35, median = 1.27, SD = 0.68). We then aggregated values at the ecoregion scale to provide meaningful biogeographical comparisons (Fig. S2). The highest values were observed in the Heard and Macdonald Islands (5.96%), the Central and Southern Great Barrier Reef (4.97%), the Ross Sea (4.90%), the Coral Sea (4.52%), and the Kerguelen Islands (3.78%; Fig. S3). Moderate values were also found in the Cortezian ecoregion (3.39%), Torres Strait Northern Great Barrier Reef (3.17%), Arnhem Coast to Gulf of Carpenteria (3.16%), Tweed-Moreton (3.10%), and Gulf of Papua (2.75%). In contrast, the lowest no-take MPA coverage was found in the Beaufort Sea - continental coast and shelf (0.34%), Chukchi Sea (0.35%), Hudson Complex (0.49%), Black Sea (0.55%), and Kamchatka Shelf and Coast (0.56%; Fig. S4). These patterns show that the top-performing ecoregions are primarily located in the coastal waters surrounding Australia (particularly in the Great Barrier Reef), the waters around the French Southern and Antarctic Lands, the Heard and McDonald Islands, and Antarctic regions, as well as various coastal waters of the Coral Triangle and Pacific tropical islands, and the southwestern Indian Ocean, while the least performing ecoregions are concentrated in Arctic waters and enclosed or semi-enclosed seas (Fig. 1 (B); Fig. S1-S3).

The geographic distribution of these protection gaps highlights systematic biases in MPA placement. While coastal waters around Australia, Antarctic regions, and remote oceanic islands showed the highest protection levels (up to 5.96% in some ecoregions), Arctic waters and enclosed seas demonstrated critically low coverage (often < 1.00%). These spatial patterns provide a meaningful background for global policy to identify areas and regions that require further efforts to protect elasmobranchs.

To evaluate whether the current spatial configuration of no-take MPAs provides better coverage for elasmobranchs than random placement would achieve, we compared observed species-level MPA coverage against a spatially explicit null model that randomised MPA placement within each country’s EEZ while maintaining the same total protected area. We found that for no-take MPAs, 63.87% of all species were under-represented, 23.93% over-represented, and 12.20% equally represented (Fig. 1 C). The most under-represented species were the Endangered Dusky Shark (*Carcharhinus obscurus*; z-score = −13.82), the Data Deficient Pencil Shark (*Hypogaleus hyugaensis*; −13.69) and the Critically Endangered Scalloped Hammerhead (*Sphyrna lewini*; −13.66; Table S2). Conversely, the mostly over-represented species were the Least Concerned Cape Sleeper Ray (*Narke capensis*; Inf), the Critically Endangered Shorttail Nurse Shark (*Pseudoginglymostoma brevicaudatum*; Inf), and the Least Concerned Viper Dogfish (*Trigonognathus kabeyai*; 32.80; Table S2).

These findings reveal that current MPA networks provide variable strategic benefit for elasmobranch conservation, with most species receiving protection levels at or below what would be expected from random placement. While MPAs were generally not designed with elasmobranchs as primary conservation targets, the fact that most under-represented species (53.37% Least Concern, 17.90% Vulnerable) mirror the overall species composition suggests opportunities exist to improve protection for threatened elasmobranchs through strategic expansion or redesign of MPA networks. This need is particularly acute in regions like the Mediterranean, where studies show that threatened elasmobranchs experience higher catch rates inside partially protected areas than in unprotected waters, indicating poor management effectiveness (Régnier et al. 2024, Sebastian et al. 2025).

To determine whether threatened species are more likely to receive worse coverage than random MPA placement, we compared the distribution of IUCN statuses between all species and those classified as under-represented. We found similar patterns between all species and those under-represented in no-take MPAs. Among them, the majority were classified as Least Concern (51.03%), followed by Vulnerable (17.90%), Near Threatened (11.21%), Endangered (11.73%), and Critically Endangered (8.13%; Fig. S5). A comparable trend was observed in under-represented species within no-take MPAs, where Least Concern species also dominated (53.37%), followed by Vulnerable (17.79%), Near Threatened (10.90%), Endangered (11.38%), and Critically Endangered (6.57%). While the relative proportions are consistent, Critically Endangered species appear slightly less represented within no-take MPAs. Overall, there is no clear evidence that threatened species (i.e., IUCN status Vulnerable, Endangered, or Critically Endangered) are disproportionately under-represented compared to other IUCN statuses (Fig. S5).

To identify whether current MPA placement performs worse or better than random allocation, we mapped mean standardized effect sizes (SES) across all species present in each grid cell and aggregated values at the ecoregion scale (Fig. S6). Mean Standardised Effect Size (SES) values were highest in Antarctic and remote oceanic island ecoregions, with the five highest values in Cocos-Keeling/Christmas Island (2.91), East Antarctic Dronning Maud Land (2.77), Ross Sea (2.77), Chagos (2.47), and Phoenix/Tokelau/Northern Cook Islands (2.30; Fig. S7). Other positive values were mostly found in tropical waters surrounding oceanic islands of the Pacific and Atlantic Oceans, as well as in waters further away from the coast (i.e., 100 nm from the coast), surrounding Africa and Central America (Fig. 1D). As opposed, mean SES values were lowest in polar and northern temperate regions, with the five most negative values found in Peter the First Island (−10.70), Amundsen/Bellingshausen Sea (−10.70), White Sea (−9.91), Weddell Sea (−8.87), and Beaufort Sea - continental coast and shelf (−8.78; Fig. S8). Additional low values were observed in northern regions, including North America, Northern Europe, and Northern East Asian regions (Fig. 1D). They were further lower in coastal waters of Australia, South American waters, coastal waters of the Coral Triangle, and Mediterranean waters. Most coastal waters of the Caribbean and Western Africa had mean SES values close to zero (Fig. 1D).

The spatial analysis of standardized effect sizes demonstrates that many of the world’s most biodiverse marine regions, including much of the Indo-Pacific, Mediterranean, and coastal Africa, show negative SES values, indicating that elasmobranch ranges in these areas overlap less with MPAs than would be expected by chance. However, range overlap analyses have inherent limitations, as they do not account for population sizes, critical habitats, migratory patterns, or life history characteristics that determine the ecological significance of protected areas for different species. Conversely, positive SES values concentrated in remote oceanic areas suggest that current MPA networks may be biased toward locations with lower human use rather than higher conservation need.

These findings echo broader concerns about MPA quality versus quantity, where recent analyses show that a quarter of assessed MPA coverage globally is not implemented, and one-third is incompatible with nature conservation due to lack of regulations or allowance of high-impact activities (Davies et al. 2017). For elasmobranchs specifically, our results suggest that achieving meaningful conservation outcomes will require both expanding coverage in priority areas and improving management effectiveness in existing protected zones.

### 3.2. Global conservation priorities

#### 3.2.1 Species-level protection outcomes

To identify priority areas for elasmobranch conservation based on IUCN threat status, FUSE, and EDGE2 dimensions, we employed the CAPTAIN algorithm. As a validation step, we first confirmed that CAPTAIN’s spatial priorities effectively targeted species according to their respective dimension scores at the 10% budget level using Beta regression models. Our results revealed significant relationships between dimension scores and species-level protection.

IUCN-based prioritisation showed a significant positive relationship between species’ threat status and their level of protection (coefficient = 0.12, SE = 0.03, z = 3.94, p < 0.001, pseudo-R² = 0.02), with species gaining an average of 2.90% additional range protection for each increase in threat category (average marginal effect = 2.90%, SE = 0.73; Fig. 2A). FUSE-based prioritisation showed a significant but weak relationship between species’ scores and their level of protection (coefficient = 0.14, SE = 0.07, z = 2.09, p = 0.04, pseudo-R² = 0.01), with species gaining an average of 3.56% additional range protection for each increase in FUSE score (average marginal effect = 3.56%, SE = 1.70; Fig. 2B). EDGE2-based prioritisation demonstrated the strongest relationship between species’ scores and their level of protection (coefficient = 0.46, SE = 0.08, z = 6.04, p < 0.001, pseudo-R² = 0.03), with species gaining an average of 11.39% additional range protection for each increase in EDGE2 score (average marginal effect = 11.39%, SE = 1.85; Fig. 2C). The very low pseudo-R² values likely reflect both the discretisation of dimensions of conservation value into only five categories and the substantial influence of species’ geographic distributions and spatial optimisation dynamics on protection outcomes. Nevertheless, the significant positive relationships confirm that CAPTAIN effectively targeted species according to their dimensions of conservation value.

**Fig. 2.**
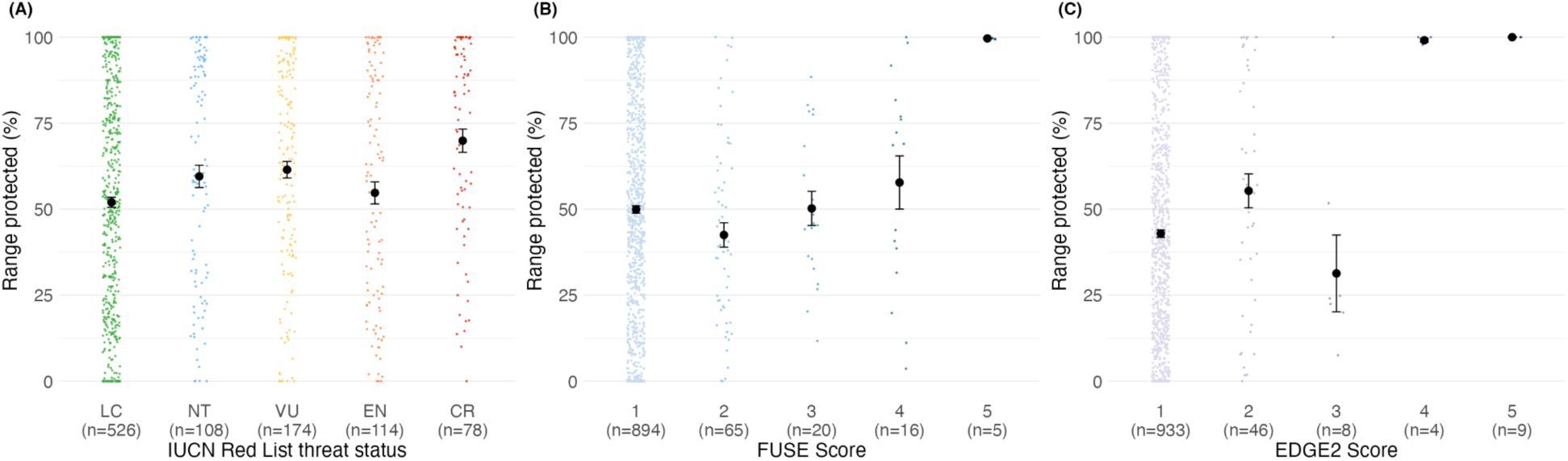
Relationship between species-level protection by CAPTAIN prioritisation and dimension of conservation value. **(A)** IUCN Red List threat status (LC = Least Concern, NT = Near Threatened, VU = Vulnerable, EN = Endangered, CR = Critically Endangered), **(B)** FUSE scores (categories 1-5 represent discretised bins of continuous scores for input into CAPTAIN), and **(C)** EDGE2 scores (categories 1-5 represent discretised bins of continuous scores for input into CAPTAIN). Black points represent mean protection values, error bars show ±1 standard error, and points show individual species (jittered for visibility); colors indicate category membership with IUCN status following Red List conventions (green for LC to red for CR), FUSE scores using a blue gradient (light to dark blue for increasing FUSE), and EDGE2 scores using a green gradient (light to dark purple for increasing EDGE2). Sample sizes for each category are shown in parentheses on the x-axis. Beta regression models revealed significant positive relationships between range protection and threat status (A; average increase of 2.87 percentage range protected per status, p < 0.001), FUSE (B; average increase of 3.50 percentage range protected per score, p = 0.04), and EDGE2 (C; average increase of 9.81 percentage range protected per score, p < 0.001).

#### 3.2.2 Geographic priority patterns

To identify global priority areas for elasmobranch conservation, we applied CAPTAIN to generate spatial priorities at the 10% budget level based on three dimensions of conservation value (IUCN status, FUSE and EDGE2). We found distinct priority patterns across dimensions, with all three concentrating priorities in coastal and continental shelf waters globally, but with notable differences in spatial distribution. Priorities based on IUCN status (Fig. 3A) showed the most dispersed pattern of high-priority cells (MNN = 51.14 km, SD = 17.18 km). The FUSE dimension (Fig. 3B) displayed intermediate spatial dispersion (MNN = 49.59 km) but the most heterogeneous distribution (SD = 26.77 km). The EDGE2 dimension (Fig. 3C) showed the most clustered pattern (MNN = 48.21 km) with the most uniform spatial distribution (SD = 8.98 km) (see Fig. 3A-C for spatial comparison).

**Fig. 3.**
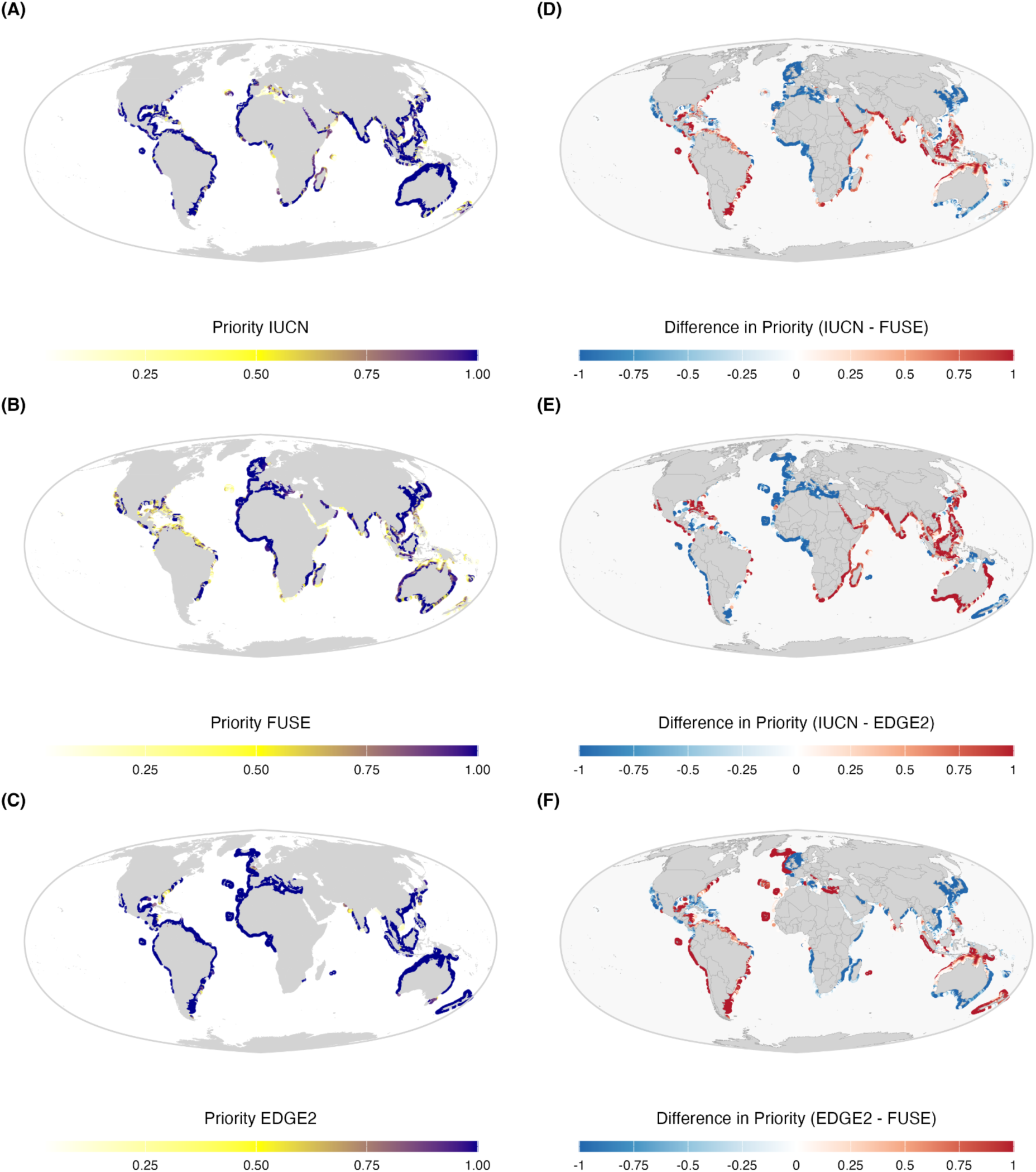
Global conservation priority maps for elasmobranchs and their pairwise comparisons. **(A)** Priority values based on IUCN status, highlighting areas with concentrations of species at highest extinction risk. **(B)** Priority values based on Functionally Unique Specialised and Endangered (FUSE) scores. **(C)** Priority values based on Evolutionary Distinct and Globally Endangered (EDGE2) scores. **(D)** Differences between IUCN and FUSE priorities. **(E)** Differences between IUCN and EDGE2 priorities. **(F)** Differences between EDGE2 and FUSE priorities. Darker colours in panels A-C indicate higher conservation priority, while darker colours in panels D-F indicate larger differences between prioritisation approaches.

We found a 46.50% congruence between areas of high priority (> 0.90) across all three indices (Fig. S9; Congruence test p-value: FUSE vs IUCN < 0.01, EDGE2 vs IUCN < 0.05 and FUSE vs EDGE2 = 0.06). Areas of congruence included coastal waters of Australia; Southeast Asian coastal regions; coastal areas of the Indian Ocean; Atlantic coastal regions of Africa; European Atlantic coastal waters; Caribbean and Eastern Pacific coastal areas; and South American Atlantic coastal waters (Fig. S9).

The substantial congruence of 46.50% across all three indices represents a remarkably high level of consensus for systematic conservation planning exercises. Studies comparing different conservation planning algorithms typically find much lower agreement, with spatial solutions often varying significantly depending on the underlying optimisation approach and target criteria (IUCN WCPA 2018). This convergence suggests that certain coastal regions, particularly around Australia, Southeast Asia, Atlantic Africa, and the Caribbean, represent robust conservation priorities regardless of whether protection is motivated by extinction risk, functional uniqueness or specialisation or evolutionary distinctiveness.

However, substantial differences between prioritisation solutions were also evident (Fig. 3; Fig. S10). Comparing IUCN with FUSE priorities (Fig. 3D), where red areas indicate higher IUCN priorities and blue areas indicate higher FUSE priorities, IUCN showed higher priorities scattered in coastal regions including parts of northern Europe, areas of the Mediterranean Sea, and various Indo-Pacific coastal waters. In contrast, FUSE showed higher priorities in much of tropical America (both Pacific and Atlantic coasts), western and central African continental waters, and parts of Southeast Asia. When comparing IUCN with EDGE2 (Fig. 3E), where red areas indicate higher IUCN priorities and blue areas indicate higher EDGE2 priorities, IUCN maintained higher priorities in similar regions as the IUCN-FUSE comparison, including Mediterranean areas and scattered coastal regions globally. EDGE2 showed distinctly higher priorities in the Australia/New Zealand region, parts of South American coastal waters, and some North American coastal areas. The direct comparison between EDGE2 and FUSE revealed complementary patterns (Fig. 3F), where red areas indicate higher EDGE2 priorities and blue areas indicate higher FUSE priorities. EDGE2 showed higher priorities particularly in Australia/New Zealand waters and southern South American coastal regions. FUSE demonstrated higher priorities in much of tropical American waters, western and central African continental shelves, the Mediterranean Sea, and parts of the Indo-Pacific region.

The application of three complementary dimensions of conservation value (IUCN status, FUSE, and EDGE2) revealed both convergent and divergent spatial priorities for elasmobranch conservation. Our use of the CAPTAIN algorithm, which employs reinforcement learning to optimise area selection, enabled us to identify spatial priorities tailored to each conservation perspective while accounting for species distributions and budget constraints. This optimisation-based approach efficiently identifies areas that maximise protection for targeted dimensions of conservation value, going beyond traditional ad-hoc or expert-driven prioritisation methods.

The distinct spatial patterns among dimensions reflect their different conservation objectives and reveal important complementarities. The methodological differences between these dimensions are not merely technical. FUSE assumes linear increases in extinction risk across IUCN statuses while EDGE2 assumes nonlinear (doubling) increases, leading to fundamentally different species rankings and spatial priorities (Castiglione et al. 2020). FUSE’s emphasis on tropical coastal ecosystems aligns with functional diversity hotspots where unique ecological roles are concentrated, while EDGE2’s focus on temperate regions like Australia and New Zealand reflects areas where evolutionarily ancient lineages persist (Pimiento et al. 2023). The IUCN approach’s more dispersed pattern captures the global distribution of immediate extinction threats.

Our findings complement broader marine conservation prioritisation efforts, which have identified that achieving representative 30% ocean protection would require targeting continental coasts, island arcs, oceanic islands, and regions like the Coral Triangle and Caribbean Sea (Lewis et al. 2017). The spatial overlap between our elasmobranch-specific priorities and these broader biodiversity patterns suggests that protecting elasmobranchs can contribute to ecosystem-wide conservation goals.

The concentration of all three prioritisations in coastal and continental shelf waters reflects both the ecological reality of elasmobranch distributions and the practical constraints of conservation implementation (Compagno 1990; Dulvy et al. 2014, 2021). Previous analyses of chondrichthyan evolutionary distinctiveness have highlighted the exceptional conservation value of this group, with the average species embodying 26 million years of unique evolutionary history, making them the most evolutionarily distinct major radiation of jawed vertebrates (Stein et al. 2018). The substantial spatial congruence, with 46.50% of high-priority areas prioritised across all three approaches, may reflect that all indices incorporate threat status through weighting (FUSE, EDGE2) or explicitly (IUCN), though it could also indicate genuine co-occurrence of multiple dimensions of conservation value. Regardless of the underlying mechanism, this convergence is ecologically meaningful and aligns with known patterns of marine biodiversity concentration: these consensus regions (waters around Australia, Southeast Asia, parts of the Indian Ocean, Atlantic Africa, European Atlantic coasts, the Caribbean and Eastern Pacific, and South American Atlantic waters) overlap substantially with recognised marine biodiversity hotspots (Ramírez et al. 2017; Pimiento et al. 2023) and represent areas where conservation action would deliver benefits across multiple dimensions of conservation value. Our spatial prioritisation translates these complementary species-level dimensions of conservation value into actionable geographic priorities.

The divergent patterns also highlight important trade-offs in conservation strategy. The spatial divergence between indices reveals where different combinations of dimensions of conservation value concentrate: IUCN status shows broader geographic coverage reflecting immediate conservation needs across all threatened species regardless of their functional or evolutionary characteristics, FUSE highlights tropical coastal ecosystems harbouring functionally unique and specialised threatened species, and EDGE2 emphasises temperate regions (particularly around Australia/New Zealand) where evolutionarily distinct lineages face elevated extinction risk. Areas prioritised by FUSE for their functional uniqueness and specialisation may require different management approaches than those identified by EDGE2 for evolutionary distinctiveness. These differences reflect the fundamental challenge in systematic conservation planning of balancing comprehensiveness (representing all dimensions of conservation value) with efficiency (maximising conservation outcomes per unit cost; Margules & Pressey 2000; Kukkala & Moilanen 2013). Our multi-dimensional approach addresses this challenge by identifying both consensus areas suitable for immediate protection and dimension-specific priorities that may require targeted management strategies.

The 10% budget constraint used in our analysis aligns with global targets for no-take MPA coverage and represents a pragmatic starting point for strategic conservation planning. However, achieving meaningful conservation outcomes requires more than simply meeting area-based targets. The high degree of spatial congruence we observed suggests that even limited conservation resources can be allocated effectively by focusing on consensus priority areas, but successful implementation will require strategic placement that accounts for local management capacity, critical habitats (such as nursery grounds and aggregation sites), connectivity between protected areas, and species-specific movement patterns and life history requirements (Kyne et al. 2023; Sacre et al. 2025). Our prioritisation framework provides a biogeographically informed foundation for such strategic placement, but must be integrated with finer-scale ecological and socioeconomic considerations during implementation.

By applying CAPTAIN across all three indices, we generated a multi-dimensional prioritisation framework that identifies areas where threat, functional uniqueness and specialisation, and evolutionary distinctiveness converge, alongside regions where specific facets of biodiversity require targeted protection. This approach ensures conservation strategies address not just the number of threatened species, but also which threatened species matter most for maintaining unique ecological functions and preserving evolutionary potential. Our prioritisation framework demonstrates the value of integrating multiple dimensions of conservation value. Rather than relying on single metrics like species richness or threat status alone, the complementary insights from threat-based, functional, and evolutionary approaches provide a more comprehensive foundation for strategic conservation decisions in an era of limited resources and conservation needs.

### 3.3. Conservation conflicts and opportunities

To identify potential conservation conflicts and opportunities, we examined the relationship between conservation priorities and commercial fishing pressure, overlaying priority values from each dimension with cumulative fishing effort from the most recent available data (2017-2020). Bivariate mapping revealed distinct spatial patterns where high conservation priorities coincide with varying levels of fishing intensity (Fig. 4). Areas shown in dark purple to red indicate regions where high priorities overlap with intensive fishing (conservation conflicts), while lighter purple areas represent high priorities with lower fishing pressure (conservation opportunities). To quantify these patterns at the ecoregion level, we aggregated cell-level data and examined mean priority values against mean fishing effort across marine ecoregions (Fig. S11). The relationship between conservation priorities and fishing pressure varied considerably across marine ecoregions and conservation indices. At the ecoregion level, most areas showed relatively low mean conservation priorities (<0.20) across all three indices, with fishing pressure spanning several orders of magnitude from 10 to 10⁵ hours per grid cell over the four-year period.

**Fig. 4.**
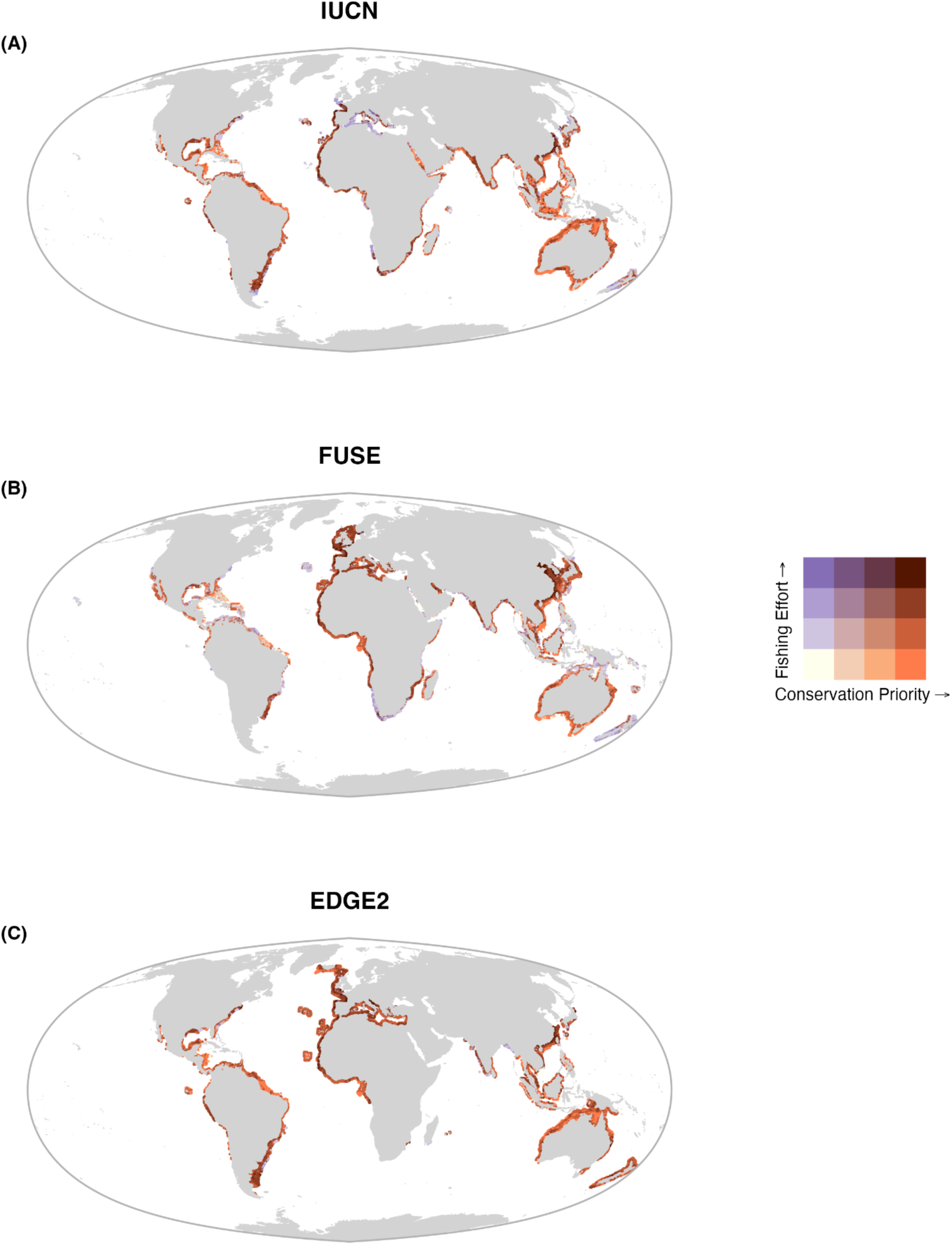
Bivariate analysis of conservation priorities and fishing pressure for elasmobranchs. Maps show the spatial relationship between conservation priority scores and fishing effort (log-transformed and normalised) across three prioritisation approaches: (A) IUCN threat-based priorities, (B) FUSE, and (C) EDGE2 priorities. Only grid cells with priority values >0 are displayed. Colours represent the intersection of conservation priority and fishing intensity, with purple tones indicating conservation opportunities (high priority, low fishing), orange/red tones indicating conservation conflicts (high priority, high fishing), and intermediate colours showing varying combinations.

Several ecoregions emerged as high-priority areas with relatively low fishing pressure, representing potential conservation opportunities. For IUCN priorities (Fig. S11A), the Torres Strait Northern Great Barrier Reef and Uruguay-Buenos Aires Shelf ecoregions showed the highest conservation values (>0.40) with moderate fishing pressure (around 100-1000 hours). The Floridian, Sunda Shelf/Java Sea, and Exmouth to Broome ecoregions also displayed elevated priorities with low to moderate fishing intensity. The Southern China and East China Sea ecoregions demonstrated moderate conservation priority scores (approximately 0.25-0.30) but experienced substantially higher fishing pressure (>10,000 hours), indicating potential conservation challenges in these heavily exploited areas.

FUSE priorities (Fig. S11B) identified a different set of high-priority ecoregions, with several areas showing conservation conflicts. The East China Sea emerged as a high-priority area (>0.50) but with very high fishing pressure (>10,000 hours over the four-year period), indicating a significant conservation conflict. The Ionian Sea, Southern Vietnam, Western Mediterranean, and Celtic Seas showed elevated conservation values (>0.40) with moderate to high fishing pressure. The Southern China, Yellow Sea, and Gulf of Tonkin ecoregions showed intermediate priorities (>0.20) but with high fishing pressure, representing additional areas of concern.

EDGE2 priorities (Fig. S11C) highlighted the Faroe Plateau as the highest-priority ecoregion (>0.70) with relatively low fishing pressure, representing a clear conservation opportunity. The Torres Strait Northern Great Barrier Reef showed high evolutionary distinctiveness with moderate fishing pressure. Several other high-priority areas, including the Exmouth to Broome, Arafura Sea, and Uruguay-Buenos Aires Shelf ecoregions (>0.40), experienced low to moderate fishing intensity.

Multiple ecoregions showed the combination of moderately high conservation priority and high to very high fishing pressure. The East China Sea, Yellow Sea, Gulf of Tonkin, and Adriatic Sea consistently appeared across indices as areas where significant fishing effort (1,000-100,000 hours over the four-year period) coincides with elevated biodiversity values, representing conservation challenges.

The juxtaposition of conservation priorities with fishing pressure patterns revealed a clear dichotomy between regions where conservation can be proactively implemented in relatively pristine areas and those requiring immediate intervention to address ongoing biodiversity losses under intensive exploitation. The most encouraging findings emerged from areas showing high conservation priority coupled with low to moderate fishing pressure, representing clear conservation opportunities. The Faroe Plateau, Torres Strait Northern Great Barrier Reef, and Uruguay-Buenos Aires Shelf exemplified such regions, where high biodiversity values coincide with manageable levels of human impact. These areas offer the potential for establishing highly protected marine protected areas that could serve as refugia for threatened species while maintaining ecosystem integrity. The relatively low fishing pressure in these regions suggests that conservation interventions could be implemented with minimal socioeconomic disruption, potentially fostering greater stakeholder acceptance and compliance.

However, our analysis also revealed conservation conflicts where high biodiversity value intersects with intensive fishing activity. The East China Sea emerged as a particularly acute example, showing high FUSE priority scores (>0.50) while experiencing fishing pressure exceeding 10,000 hours. A pattern that mirrors broader global trends where intensive fishing pressure, including from artisanal and subsistence fishing in remote regions, can cause serious depletion of large coastal elasmobranchs, with most elasmobranch stocks having collapsed due to unsustainable fishing compounded by their low productivity (Davidson and Dulvy 2017; Dulvy et al. 2017).

The persistence of moderate to high conservation priorities in heavily exploited regions like the Yellow Sea, Gulf of Tonkin, and Adriatic Sea highlights the urgent need for innovative management approaches. These areas represent classic examples of conflicts between marine nature conservation and fishery interests, where there is often a glaring lack of dialogue between stakeholders representing these two interests. The challenge is compounded by the fact that elasmobranch populations have declined by 80% or more in many regions across the globe, predominantly due to unsustainable fisheries driven by high demand for fins and meat, together with high levels of bycatch (Kriegl et al. 2021).

The temporal dimension of these conflicts deserves particular attention. Recent research using machine learning to understand how fishing fleets respond to marine protected areas reveals that fishing effort changes in complex ways, with decreases occurring both inside and outside new MPAs through changes in fish stocks, costs, and profitability (Mcdonald et al. 2024). This suggests that conservation interventions in high-conflict areas may have cascading effects that extend beyond immediate MPA boundaries.

The geographic clustering of conservation conflicts in Asian seas (East China Sea, Yellow Sea, Gulf of Tonkin) and the Mediterranean reflects broader regional patterns of intensive coastal development and fisheries exploitation. In regions like the Bay of Bengal, increasing global pressure from fishing and trade on cartilaginous fish is pushing stocks close to critical limits, with high fishing pressure from both targeted and bycatch fisheries impacting populations tremendously (Dulvy et al. 2008). These regional hotspots require coordinated international management approaches that address both local fishing pressures and broader market-driven exploitation.

The contrast between opportunity and conflict areas also highlights important lessons for marine spatial planning. The effectiveness of MPAs depends significantly on their level of protection, with highly and fully protected MPAs providing substantially greater conservation benefits than lightly protected areas that allow commercial fishing. In opportunity areas, this suggests investing in high-protection MPAs, while conflict areas may require innovative approaches such as seasonal closures, gear restrictions, or transitional management strategies.

Our analysis reveals that effective elasmobranch conservation will require a dual strategy: proactive protection of high-value, low-conflict areas combined with intensive management intervention in high-conflict regions. Given the ecological and demographic diversity of elasmobranchs, a range of fisheries management measures are generally preferable to ‘one size fits all’ conservation actions, with some species requiring strict protections to minimise mortality while others can sustain fishing under effective catch limits (Sève et al. 2023). Success will ultimately depend on developing context-specific solutions that balance conservation imperatives with socioeconomic realities in each region.

### 4.4. Study limitations

While our analysis provides valuable insights into global elasmobranch conservation priorities, several methodological limitations warrant acknowledgment.

Our reliance on the World Database on Protected Areas (WDPA), while representing the most comprehensive global MPA dataset available, suffers from significant reporting gaps and inconsistencies. Many jurisdictions fail to report newly established MPAs, particularly smaller, locally managed areas and community-based conservation initiatives (Mouton et al. 2025; Farhadinia et al. 2022). Recent satellite analyses reveal that industrial fishing occurs in 47% of coastal MPAs worldwide, with two-thirds of vessel detections untracked by public monitoring systems (Seguin et al. 2025), suggesting our gap analysis may have underestimated actual fishing pressure while overestimating effective protection levels.

Our global-scale prioritisation inherently produces broad priority areas that cannot be directly designated as individual MPAs. The priority regions identified represent extensive ocean areas requiring subdivision into practical management units and should be interpreted as regions where networks of complementary MPAs need strategic establishment rather than specific MPA boundaries. Additionally, the 10% budget constraint, while realistic given current funding limitations, may be insufficient given that 97.4% of marine species have less than 10% of their range in existing protected areas (Holness et al. 2022). Finally, our fishing pressure analysis captures industrial fishing comprehensively but has limited coverage of small-scale and artisanal fisheries (Kroodsma et al. 2018, Paolo et al. 2024), particularly in regions like the Caribbean, Mediterranean, and Coral Triangle, potentially underestimating total fishing pressure in those areas.

Despite these limitations, our analysis represents the first global assessment integrating threat status, functional uniqueness, and evolutionary distinctiveness for elasmobranch conservation prioritisation using the CAPTAIN reinforcement learning algorithm, providing a robust foundation for strategic conservation planning

## 5. Conclusion

Our comprehensive global analysis reveals critical gaps in current elasmobranch protection while providing a robust framework for strategic conservation planning. The stark finding that no-take marine protected areas cover only 3% of species ranges on average, with 64% of species under-represented relative to random placement, underscores the urgent need for targeted conservation action. However, our multi-dimensional prioritisation approach offers hope by identifying consensus priority regions where threat-based, functional, and evolutionary perspectives align, providing clear targets for conservation investment. The 46% spatial congruence across all three indices demonstrates that despite methodological differences, certain coastal regions, particularly around Australia, Southeast Asia, Atlantic Africa, and the Caribbean, represent robust conservation priorities regardless of the specific dimensions of conservation value employed. While our analysis reveals conservation conflicts in heavily exploited regions like the East China Sea and Mediterranean, it also identifies promising opportunities in areas like the Faroe Plateau and Torres Strait where high conservation value coincides with manageable fishing pressure. Moving forward, protecting elasmobranchs will require a dual strategy: proactive establishment of highly protected marine reserves in low-conflict priority areas, coupled with intensive management interventions in high-conflict regions where significant biodiversity value intersects with intensive human use. Success will ultimately depend on translating these global priority insights into practical, locally appropriate conservation actions that balance ecological imperatives with socioeconomic realities. As the world mobilises to meet the 30×30 target, our findings provide an evidence-based foundation for ensuring that elasmobranchs, among the ocean’s most threatened yet ecologically vital species, are adequately represented in the global expansion of marine protection.

## 7 Acknowledgements

The authors gratefully acknowledge David Mouillot, Camille Albouy, Laure Velez, and John F. Griffin for their assistance with data curation and for insightful discussions that advanced the development of this manuscript.

## 9 Data and materials availability

All data and code used to support the findings of this study are publicly available at: https://github.com/TheophileMt92/CAPTAIN_elasmo_manuscript.

## References

1. Albano, Patricia S, Chris Fallows, Monique Fallows, Olivia Schuitema, Anthony TF Bernard, Oliver Sedgwick, and Neil Hammerschlag. 2021. “Successful parks for sharks: No-take marine reserve provides conservation benefits to endemic and threatened sharks off South Africa.” Biological conservation 261: 109302.

2. Albouy, Camille, Valentine L Delattre, Bastien Mérigot, Christine N Meynard, and Fabien Leprieur. 2017. “Multifaceted biodiversity hotspots of marine mammals for conservation priorities.” Diversity and Distributions 23 (6): 615–626.

3. Aminian-Biquet, Juliette, Jennifer Sletten, Timothé Vincent, Margherita Pieraccini, Betty Queffelec, Anastasiya Laznya, Natașa Vaidianu, Joachim Claudet, Juliette Young, and Barbara Horta e Costa. 2025. “Major data gaps and recommendations in monitoring regulations of activities in EU marine protected areas.” npj Ocean Sustainability 4 (1): 3.

4. Baddeley, Adrian, and Rolf Turner. 2014. “Package ‘spatstat’.” The Comprehensive R Archive Network () 146.

5. Ball, Ian R, Hugh P Possingham, and Matthew Watts. 2009. “Marxan and relatives: software for spatial conservation prioritisation.” Spatial conservation prioritisation: Quantitative methods and computational tools 14: 185–196.

6. Baum, Julia K, and Boris Worm. 2009. “Cascading top-down effects of changing oceanic predator abundances.” Journal of animal ecology 78 (4): 699–714.

7. Bond, Mark E, Jasmine Valentin-Albanese, Elizabeth A Babcock, Debra Abercrombie, Norlan F Lamb, Ashbert Miranda, Ellen K Pikitch, and Demian D Chapman. 2017. “Abundance and size structure of a reef shark population within a marine reserve has remained stable for more than a decade.” Marine Ecology Progress Series 576: 1–10.

8. Breiman, Leo, Adele Cutler, Andy Liaw, Matthew Wiener, and Maintainer Andy Liaw. 2018. “Package ‘randomforest’.” University of California, Berkeley: Berkeley, CA, USA 81: 1–29.

9. Cade, Brian S, and Barry R Noon. 2003. “A gentle introduction to quantile regression for ecologists.” Frontiers in Ecology and the Environment 1(8): 412–420.

10. Convention on Biological Diversity (CBD). 2022. Kunming-Montreal Global biodiversity framework Fifteenth meeting.

11. Compagno, Leonard JV. 1990. “Alternative life-history styles of cartilaginous fishes in time and space.” Environmental Biology of Fishes 28 (1): 33–75.

12. Davidson, Lindsay NK, and Nicholas K Dulvy. 2017. “Global marine protected areas to prevent extinctions.” Nature ecology & evolution 1 (2): 1–6.

13. Davies, TE, SM Maxwell, K Kaschner, Cristina Garilao, and Natalie C Ban. 2017. “Large marine protected areas represent biodiversity now and under climate change.” Scientific Reports 7 (1): 1–7.

14. Day, J., N. Dudley, M. Hockings, G. Holmes, D. Laffoley, S. Stolton, S. Wells, and L. Wenzel. 2019. Guidelines for applying the IUCN protected area management categories to marine protected areas. IUCN (Gland, Switzerland).

15. Dedman, Simon, Jerry H Moxley, Yannis P Papastamatiou, Matias Braccini, Jennifer E Caselle, Demian D Chapman, Joshua Eli Cinner, Erin M Dillon, Nicholas K Dulvy, and Ruth Elizabeth Dunn. 2024. “Ecological roles and importance of sharks in the Anthropocene Ocean.” Science 385 (6708): adl2362.

16. Di Lorenzo, Manfredi, Paolo Guidetti, Antonio Di Franco, Antonio Calò, and Joachim Claudet. 2020. “Assessing spillover from marine protected areas and its drivers: A meta-analytical approach.” Fish and Fisheries 21 (5): 906–915.

17. Dulvy, Nicholas K, Julia K Baum, Shelley Clarke, Leonard JV Compagno, Enric Cortés, Andrés Domingo, Sonja Fordham, Sarah Fowler, Malcolm P Francis, and Claudine Gibson. 2008. “You can swim but you can’t hide: the global status and conservation of oceanic pelagic sharks and rays.” Aquatic conservation: marine and freshwater ecosystems 18 (5): 459–482.

18. Dulvy, Nicholas K, Nathan Pacoureau, Jay H Matsushiba, Helen F Yan, Wade J VanderWright, Cassandra L Rigby, Brittany Finucci, C Samantha Sherman, Rima W Jabado, and John K Carlson. 2024. “Ecological erosion and expanding extinction risk of sharks and rays.” Science 386 (6726): eadn1477.

19. Dulvy, Nicholas K, Nathan Pacoureau, Cassandra L Rigby, Riley A Pollom, Rima W Jabado, David A Ebert, Brittany Finucci, Caroline M Pollock, Jessica Cheok, and Danielle H Derrick. 2021. “Overfishing drives over one-third of all sharks and rays toward a global extinction crisis.” Current Biology 31 (21): 4773–4787. E8.

20. Dulvy, Nicholas K, Colin A Simpfendorfer, Lindsay NK Davidson, Sonja V Fordham, Amie Bräutigam, Glenn Sant, and David J Welch. 2017. “Challenges and priorities in shark and ray conservation.” Current Biology 27 (11): R565–R572.

21. Dwyer, Ross G, Hamish A Campbell, Richard D Pillans, Matthew E Watts, Barry J Lyon, Siddeswara M Guru, Minh N Dinh, Hugh P Possingham, and Craig E Franklin. 2019. “Using individual-based movement information to identify spatial conservation priorities for mobile species.” Conservation Biology 33 (6): 1426–1437.

22. Dwyer, Ross G, Nils C Krueck, Vinay Udyawer, Michelle R Heupel, Demian Chapman, Harold L Pratt, Ricardo Garla, and Colin A Simpfendorfer. 2020. “Individual and population benefits of marine reserves for reef sharks.” Current Biology 30 (3): 480–489. E5.

23. Farhadinia, Mohammad S, Anthony Waldron, Żaneta Kaszta, Ehab Eid, Alice Hughes, Hüseyin Ambarlı, Hadi Al-Hikmani, Bayarbaatar Buuveibaatar, Mariya A Gritsina, and Iding Haidir. 2022. “Current trends suggest most Asian countries are unlikely to meet future biodiversity targets on protected areas.” Communications Biology 5 (1): 1221.

24. Ferrari, Silvia, and Francisco Cribari-Neto. 2004. “Beta regression for modelling rates and proportions.” Journal of applied statistics 31 (7): 799–815.

25. Fox, John, and Sanford Weisberg. 2018. An R companion to applied regression. Sage publications.

26. Gouhier, Tarik C, and Pradeep Pillai. 2020. “Avoiding conceptual and mathematical pitfalls when developing indices to inform conservation.” Frontiers in Ecology and Evolution 8: 263.

27. Gumbs, Rikki, Claudia L Gray, Monika Böhm, Ian J Burfield, Olivia R Couchman, Daniel P Faith, Félix Forest, Michael Hoffmann, Nick JB Isaac, and Walter Jetz. 2023. “The EDGE2 protocol: Advancing the prioritisation of Evolutionarily Distinct and Globally Endangered species for practical conservation action.” PLoS Biology 21 (2): e3001991.

28. Hanson, Jeffrey O, Richard Schuster, Matthew Strimas-Mackey, Nina Morrell, Brandon PM Edwards, Peter Arcese, Joseph R Bennett, and Hugh P Possingham. 2024. “Systematic conservation prioritization with the prioritizr R package.” Conservation Biology: e14376.

29. Heithaus, Michael R, Alejandro Frid, Aaron J Wirsing, and Boris Worm. 2008. “Predicting ecological consequences of marine top predator declines.” Trends in ecology & evolution 23 (4): 202–210.

30. Holness, Stephen D, Linda R Harris, Russell Chalmers, Deidre De Vos, Victoria Goodall, Hannah Truter, Ané Oosthuizen, Anthony TF Bernard, Paul D Cowley, and Charlene da Silva. 2022. “Using systematic conservation planning to align priority areas for biodiversity and nature-based activities in marine spatial planning: a real-world application in contested marine space.” Biological Conservation 271: 109574.

31. Isaac, Nick JB, Samuel T Turvey, Ben Collen, Carly Waterman, and Jonathan EM Baillie. 2007. “Mammals on the EDGE: conservation priorities based on threat and phylogeny.” PloS one 2 (3): e296.

32. Jacquemont, Juliette, Robert Blasiak, Chloé Le Cam, Maël Le Gouellec, and Joachim Claudet. 2022. “Ocean conservation boosts climate change mitigation and adaptation.” One Earth 5 (10): 1126–1138.

33. Jacquemont, Juliette, Charles Loiseau, Luke Tornabene, and Joachim Claudet. 2024. “3D ocean assessments reveal that fisheries reach deep but marine protection remains shallow.” Nature Communications 15 (1): 4027.

34. Jonckheere, A.R. 1954. “A distribution-free k-sample test against ordered alternatives.” Biometrika 41(1-2): 133–145.

35. Klimley, A Peter, Randall Arauz, Sandra Bessudo, Elpis J Chávez, Nicole Chinacalle, Eduardo Espinoza, Jonathan Green, Alex R Hearn, Mauricio E Hoyos-Padilla, and Elena Nalesso. 2022. “Studies of the movement ecology of sharks justify the existence and expansion of marine protected areas in the Eastern Pacific Ocean.” Environmental Biology of Fishes 105 (12): 2133–2153.

36. Koenker, R. 2005. “Quantile regression.” Cambridge University Press.

37. Koenker, R. 2024. quantreg: Quantile regression. https://CRAN.R-project.org/package=quantreg.

38. Komsta, Lukasz, and Frederick Novomestky. 2015. “Moments, cumulants, skewness, kurtosis and related tests.” R package version 14 (1).

39. Kriegl, Michael, Xochitl E Elías Ilosvay, Christian von Dorrien, and Daniel Oesterwind. 2021. “Marine protected areas: at the crossroads of nature conservation and fisheries management.” Frontiers in Marine Science 8: 676264.

40. Kroodsma, David A, Juan Mayorga, Timothy Hochberg, Nathan A Miller, Kristina Boerder, Francesco Ferretti, Alex Wilson, Bjorn Bergman, Timothy D White, and Barbara A Block. 2018. “Tracking the global footprint of fisheries.” Science 359 (6378): 904–908.

41. Kukkala, Aija S, and Atte Moilanen. 2013. “Core concepts of spatial prioritisation in systematic conservation planning.” Biological Reviews 88 (2): 443–464.

42. Kyne, Peter M, Giuseppe Notarbartolo di Sciara, Amanda Batlle Morera, Ryan Charles, Emiliano García Rodríguez, Daniel Fernando, Adriana Gonzalez Pestana, Mark Priest, and Rima W Jabado. 2023. “Important Shark and Ray Areas: a new tool to optimize spatial planning for sharks.” Oryx 57 (2): 146–147.

43. Leeper, Thomas J, Jeffrey Arnold, Vincent Arel-Bundock, and Jacob A Long. 2021. Package ‘margins’.

44. Lewis, Nai’a, Jon C Day, ’Aulani Wilhelm, Daniel Wagner, Carlos Gaymer, John Parks, Alan Friedlander, Susan White, Charles Sheppard, and Mark Spalding. 2017. “Large-scale marine protected areas: guidelines for design and management.”

45. Margules, Christopher Robert, and Robert L Pressey. 2000. “Systematic conservation planning.” Nature 405 (6783): 243–253.

46. McDonald, Gavin, Jennifer Bone, Christopher Costello, Gabriel Englander, and Jennifer Raynor. 2024. “Global expansion of marine protected areas and the redistribution of fishing effort.” Proceedings of the National Academy of Sciences 121 (29): e2400592121.

47. Medoff, Sarah, John Lynham, and Jennifer Raynor. 2022. “Spillover benefits from the world’s largest fully protected MPA.” Science 378 (6617): 313–316.

48. Moilanen, Atte, Heini Kujala, and John R Leathwick. 2009. “The Zonation framework and software for conservation prioritization.” Spatial conservation prioritization 135: 196–210.

49. Mouillot, David, David R Bellwood, Christopher Baraloto, Jerome Chave, Rene Galzin, Mireille Harmelin-Vivien, Michel Kulbicki, Sebastien Lavergne, Sandra Lavorel, and Nicolas Mouquet. 2013. “Rare species support vulnerable functions in high-diversity ecosystems.” PLoS biology 11 (5).

50. Mouton, Théophile L, Adriana Gonzalez-Pestana, Christoph A Rohner, Ryan Charles, Emiliano García-Rodríguez, Peter M Kyne, Amanda Batlle-Morera, Giuseppe Notarbartolo di Sciara, Asia O Armstrong, and Enzo Acuña. 2025. “Shortfalls in the protection of Important Shark and Ray Areas undermine shark conservation efforts in the Central and South American Pacific.” Marine Policy 171: 106448.

51. Paolo, Fernando S, David Kroodsma, Jennifer Raynor, Tim Hochberg, Pete Davis, Jesse Cleary, Luca Marsaglia, Sara Orofino, Christian Thomas, and Patrick Halpin. 2024. “Satellite mapping reveals extensive industrial activity at sea.” Nature 625 (7993): 85–91.

52. Pavoine, Sandrine, Jeanne Vallet, Anne-Béatrice Dufour, Sophie Gachet, and Hervé Daniel. 2009. “On the challenge of treating various types of variables: application for improving the measurement of functional diversity.” Oikos 118 (3): 391–402.

53. Petchey, Owen L, and Kevin J Gaston. 2006. “Functional diversity: back to basics and looking forward.” Ecology letters 9 (6): 741–758.

54. Pimiento, C, F Leprieur, D Silvestro, JS Lefcheck, Camille Albouy, DB Rasher, M Davis, J-C Svenning, and JN Griffin. 2020. “Functional diversity of marine megafauna in the Anthropocene.” Science Advances 6 (16): eaay7650.

55. Pimiento, Catalina, Camille Albouy, Daniele Silvestro, Théophile L Mouton, Laure Velez, David Mouillot, Aaron B Judah, John N Griffin, and Fabien Leprieur. 2023. “Functional diversity of sharks and rays is highly vulnerable and supported by unique species and locations worldwide.” Nature Communications 14 (1): 7691.

56. Pollock, Laura J, Wilfried Thuiller, and Walter Jetz. 2017. “Large conservation gains possible for global biodiversity facets.” Nature 546 (7656): 141–144.

57. Ramírez, Francisco, Isabel Afán, Lloyd S Davis, and André Chiaradia. 2017. “Climate impacts on global hot spots of marine biodiversity.” Science Advances 3 (2): e1601198.

58. Régnier, Thomas, Jane Dodd, Steven Benjamins, Fiona M Gibb, and Peter J Wright. 2024. “Spatial management measures benefit the critically endangered flapper skate, Dipturus intermedius.” Aquatic Conservation: Marine and Freshwater Ecosystems 34 (4): e4150.

59. Roberts, Callum M, Bethan C O’Leary, Douglas J McCauley, Philippe Maurice Cury, Carlos M Duarte, Jane Lubchenco, Daniel Pauly, Andrea Sáenz-Arroyo, Ussif Rashid Sumaila, and Rod W Wilson. 2017. “Marine reserves can mitigate and promote adaptation to climate change.” Proceedings of the National Academy of Sciences 114 (24): 6167–6175.

60. Sacre, Edmond, Ulf Bergström, and Charlotte Berkström. 2025. “Identifying priority areas for conservation to promote connectivity and mitigate the impacts of anthropogenic disturbance.” Conservation Biology: e70083.

61. Sebastian, Pascal, Serena J Stean, Ahmad Ilham Rabbani Erawan, Rinaldi Gotama, I Nengah Swarya, Lauren Dawn Sparks, Rahmadi Prasetijo, and Andhika P Prasetyo. 2025. “Quantifying the influence of environmental factors on elasmobranch distribution and abundance in a high-use marine protected area.” Marine Environmental Research: 107317.

62. Seguin, Raphael, Frédéric Le Manach, Rodolphe Devillers, Laure Velez, and David Mouillot. 2025. “Global patterns and drivers of untracked industrial fishing in coastal marine protected areas.” Science 389 (6758): 396–401.

63. Sève, Charlotte, Mokrane Belharet, Paco Melià, Antonio Di Franco, Antonio Calò, and Joachim Claudet. 2023. “Fisheries outcomes of marine protected area networks: Levels of protection, connectivity, and time matter.” Conservation Letters 16 (6): e12983.

64. Signorell, Aea, K Aho, A Alfons, N Anderegg, T Aragon, C Arachchige, A Arppe, A Baddeley, K Barton, and B Bolker. 2024. DescTools: Tools for descriptive statistics. R package version 0.99. 55. https://CRAN.R-project.org/package=DescTools

65. Silvestro, Daniele, Stefano Goria, Ben Groom, Piotr Jacobsson, Thomas Sterner, and Alexandre Antonelli. 2025. “Using artificial intelligence to optimize ecological restoration for climate and biodiversity.” bioRxiv: 2025.01. 31.635975.

66. Silvestro, Daniele, Stefano Goria, Thomas Sterner, and Alexandre Antonelli. 2022. “Improving biodiversity protection through artificial intelligence.” Nature sustainability 5 (4): 415–424.

67. Smithson, Michael, and Jay Verkuilen. 2006. “A better lemon squeezer? Maximum-likelihood regression with beta-distributed dependent variables.” Psychological methods 11 (1): 54.

68. Speed, Conrad W, Mike Cappo, and Mark G Meekan. 2018. “Evidence for rapid recovery of shark populations within a coral reef marine protected area.” Biological Conservation 220: 308–319.

69. Stein, R William, Christopher G Mull, Tyler S Kuhn, Neil C Aschliman, Lindsay NK Davidson, Jeffrey B Joy, Gordon J Smith, Nicholas K Dulvy, and Arne O Mooers. 2018. “Global priorities for conserving the evolutionary history of sharks, rays and chimaeras.” Nature ecology & evolution 2 (2): 288–298.

70. Terpstra, T.J. 1952. “The asymptotic normality and consistency of Kendall’s test against trend, when ties are present in one ranking.” Indagationes Mathematicae 14: 327–333.

71. Thuiller, Wilfried, Luigi Maiorano, Florent Mazel, François Guilhaumon, Gentile Francesco Ficetola, Sébastien Lavergne, Julien Renaud, Cristina Roquet, and David Mouillot. 2015. “Conserving the functional and phylogenetic trees of life of European tetrapods.” Philosophical Transactions of the Royal Society B: Biological Sciences 370 (1662): 20140005.

72. Watson, James EM, Ruben Venegas-Li, Hedley Grantham, Nigel Dudley, Sue Stolton, Madhu Rao, Stephen Woodley, Marc Hockings, Karl Burkart, and Jeremy S Simmonds. 2023. “Priorities for protected area expansion so nations can meet their Kunming-Montreal Global Biodiversity Framework commitments.” Integrative Conservation 2 (3): 140–155.

73. WCPA IUCN. 2018. Applying IUCN’s Global Conservation Standards to Marine Protected Areas (MPA). Delivering effective conservation action through MPAs, to secure ocean health & sustainable development. (Gland, Switzerland).

74. Welch, Heather, Tyler Clavelle, Timothy D White, Megan A Cimino, Jennifer Van Osdel, Timothy Hochberg, David Kroodsma, and Elliott L Hazen. 2022. “Hot spots of unseen fishing vessels.” Science Advances 8 (44): eabq2109.

75. Yan, Helen F, Peter M Kyne, Rima W Jabado, Ruth H Leeney, Lindsay NK Davidson, Danielle H Derrick, Brittany Finucci, Robert P Freckleton, Sonja V Fordham, and Nicholas K Dulvy. 2021. “Overfishing and habitat loss drive range contraction of iconic marine fishes to near extinction.” Science Advances 7 (7): eabb6026.

76. Zeileis, Achim, Francisco Cribari-Neto, Bettina Gruen, Ioannis Kosmidis, Alexandre B Simas, Andrea V Rocha, and Maintainer Achim Zeileis. 2016. “Package ‘betareg’.” R package 3 (2): 51.

