## Supplementary materials for "Global gaps and priorities for shark and ray conservation: Integrating threat, function, and evolutionary distinctiveness"

### Table of Contents

***Supplementary materials*** ... **2**

Table S1. Quantile regression results examining the relationship between IUCN threat status and current MPA coverage across the distribution of protection levels. ... 2

Fig S1. Distribution of current MPA coverage by IUCN threat status and quantile regression analysis. ... 3

Fig S2. Global distribution of mean percentage range in no-take marine protected areas by marine ecoregion. ... 4

Fig S3. Top 20 marine ecoregions ranked by mean percentage range overlapped by no-take marine protected areas. ... 5

Fig S4. Bottom 20 marine ecoregions ranked by mean percentage range overlapped by no-take marine protected areas. ... 6

Table S2. Top 10 elasmobranch species with the greatest over- and under-representation in no-take Marine Protected Areas relative to null model expectations. ... 7

Fig S5. IUCN threat status distribution of elasmobranch species under-represented in no-take MPAs compared to all studied species. ... 8

Fig S6. Standardized effect size (SES) of observed versus null no-take marine protected area coverage by marine ecoregion. ... 9

Fig S7. Top 20 marine ecoregions with highest mean standardized effect sizes (SES) for no-take marine protected area coverage. ... 10

Fig S8. Bottom 20 marine ecoregions with lowest mean standardized effect sizes (SES) for no-take marine protected area coverage. ... 11

Fig S9. Spatial congruence of high-priority conservation areas across three dimensions of conservation values. ... 12

Fig S10. Global spatial congruence of elasmobranch conservation priorities across three dimensions of conservation value. ... 13

Fig S11. Relationship between fishing pressure and conservation priority by marine ecoregion. ... 14

***Supplementary methods*** ... **15**

**SM1. Predicting fishing activity using random forest modelling** ... **15**

Fig SM1. Global distribution of industrial fishing effort detected by Automatic Identification System (AIS). ... 16

Fig SM2. Global distribution of fishing vessel detections using Sentinel-1 Synthetic Aperture Radar (SAR). ... 17

Fig SM3. Spatial comparison of fishing activity detected by AIS and SAR monitoring systems (2017-2020). ... 18

Table SM2. Distribution of fishing activity detection across monitoring technologies. ... 19

**SM2. Random forest model performance** ... **20**

Table SM3. Performance comparison of random forest models with different data transformations for predicting fishing effort. ... 21

Table SM4. Performance metrics of the selected log-transformed fishing hours random forest model. ... 22

Table SM5. Variable importance metrics for the log-transformed fishing hours random forest model. ... 23

Fig SM4. Integrated global fishing effort map combining AIS data and random forest model predictions from SAR data (2017-2020). ... 24

### Supplementary materials

**Table S1. Quantile regression results examining the relationship between IUCN threat status and current MPA coverage across the distribution of protection levels.** Each row represents a different quantile (τ) of the coverage distribution, with the slope coefficient indicating the change in MPA coverage (%) per unit increase in threat category (from LC=1 to CR=5). Negative slopes indicate declining coverage with increasing threat status. Standard errors (SE) were estimated using bootstrap resampling with 500 replications. The t-statistic and p-value test whether the slope differs significantly from zero at each quantile. Quantiles marked as "Yes*" indicate statistically significant relationships ($p < 0.05$), while "No" indicates non-significant relationships.

Quantile regression results for current no-take MPA coverage by IUCN threat status

| Quantile | τ | Slope | SE | t | p-value | Significant |
|---|---|---|---|---|---|---|
| **Q30** | 0.3 | 0.00 | 0.01 | 0.00 | 1.00 | No |
| **Q40** | 0.4 | -0.01 | 0.06 | -0.15 | 0.88 | No |
| **Q50** | 0.5 | -0.03 | 0.09 | -0.37 | 0.71 | No |
| **Q60** | 0.6 | -0.14 | 0.09 | -1.54 | 0.12 | No |
| **Q70** | 0.7 | -0.39 | 0.09 | -4.17 | 0.00 | Yes* |
| **Q80** | 0.8 | -0.77 | 0.12 | -6.44 | 0.00 | Yes* |
| **Q90** | 0.9 | -1.38 | 0.25 | -5.43 | 0.00 | Yes* |

*Note:*

Bootstrap standard errors with 500 replications. Significant quantiles ($p < 0.05$) are highlighted.

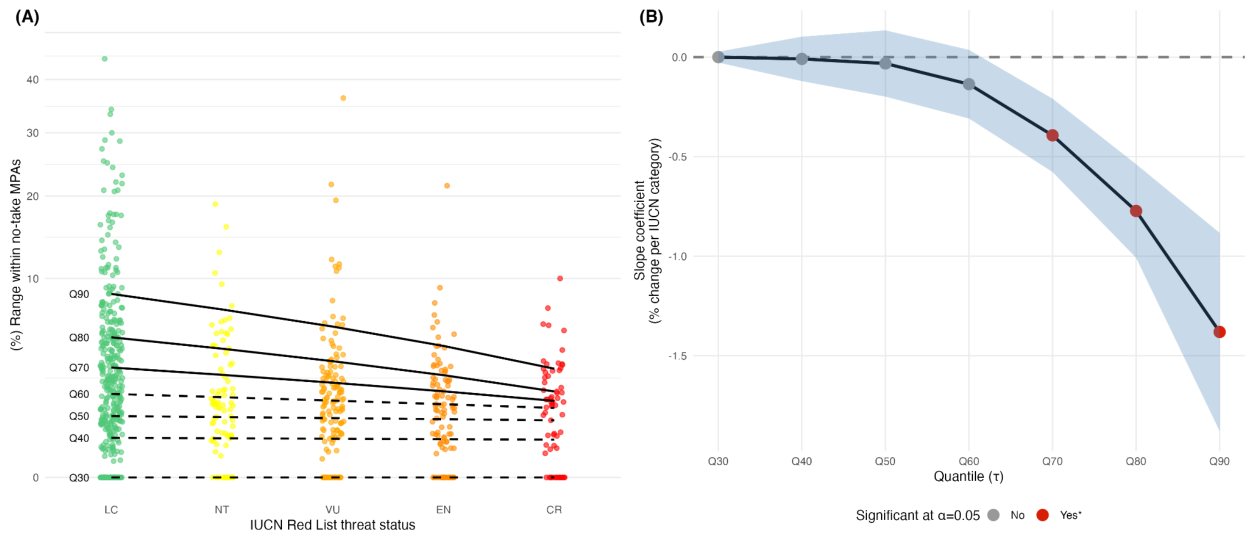


**Fig S1. Distribution of current MPA coverage by IUCN threat status and quantile regression analysis. (A)** Distribution of no-take MPA coverage across IUCN Red List threat categories, showing individual species values (colored points) and fitted quantile regression lines (Q30-Q90). Solid lines indicate statistically significant relationships ($p < 0.05$; Q70-Q90), while dashed lines indicate non-significant relationships (Q30-Q60). **(B)** Slope coefficients from quantile regression models across the distribution. Points represent slope estimates at each quantile, and shaded regions represent 95% confidence intervals around each estimate (based on bootstrap standard errors).

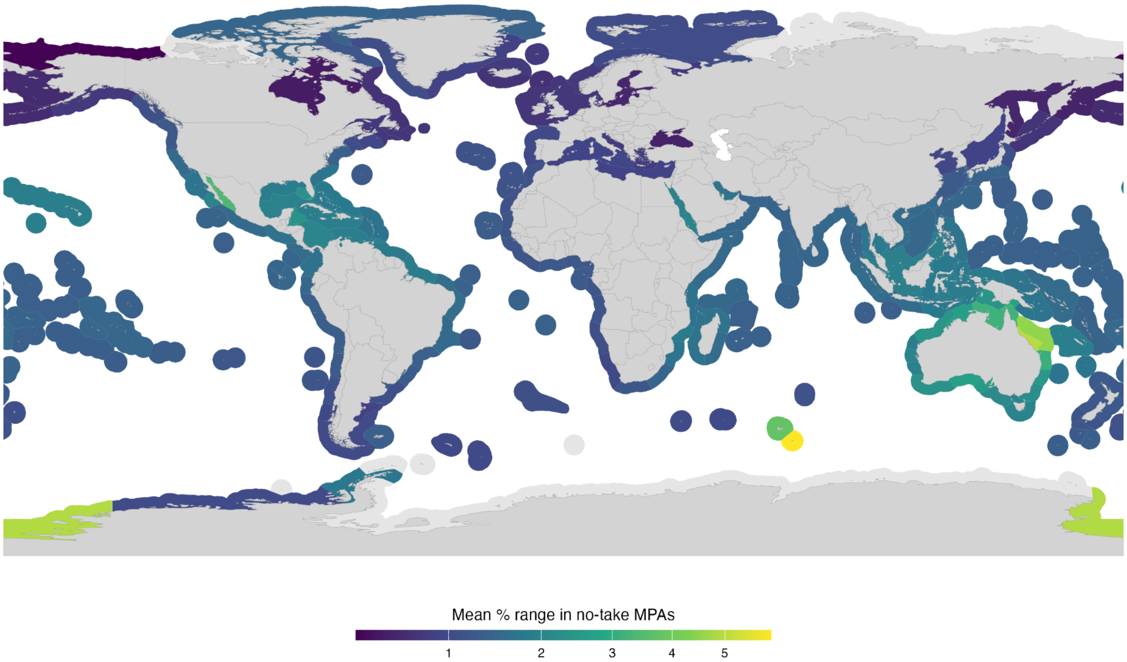


**Fig S2. Global distribution of mean percentage range in no-take marine protected areas by marine ecoregion.** The map displays the mean percentage of species ranges within no-take marine protected areas (MPAs) across the world's marine ecoregions, based on the Marine Ecoregions of the World (MEOW) classification system. Colour intensity represents the level of no-take MPA coverage, with darker blue indicating lower mean coverage percentages (square-root transformed scale). Light grey areas indicate ecoregions with no species data. Land areas are shown in darker grey for reference. The analysis reveals substantial variation in conservation coverage across different marine biogeographic regions, with some ecoregions achieving notably higher levels of no-take protection than others.

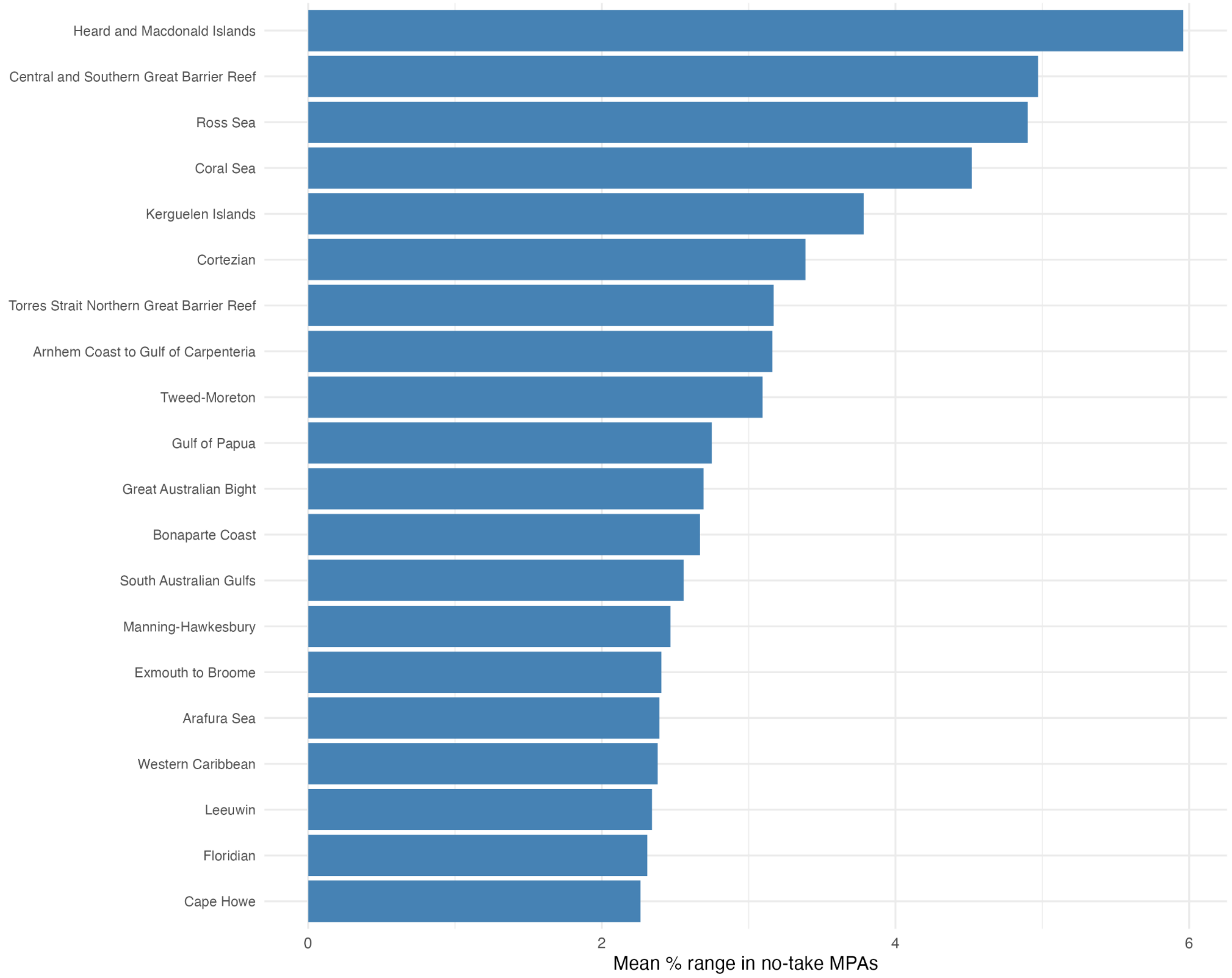


**Fig S3. Top 20 marine ecoregions ranked by mean percentage range overlapped by no-take marine protected areas.** Bar chart showing the marine ecoregions with the highest mean percentage of species ranges protected within no-take marine protected areas. Ecoregions are ranked in ascending order of mean coverage.

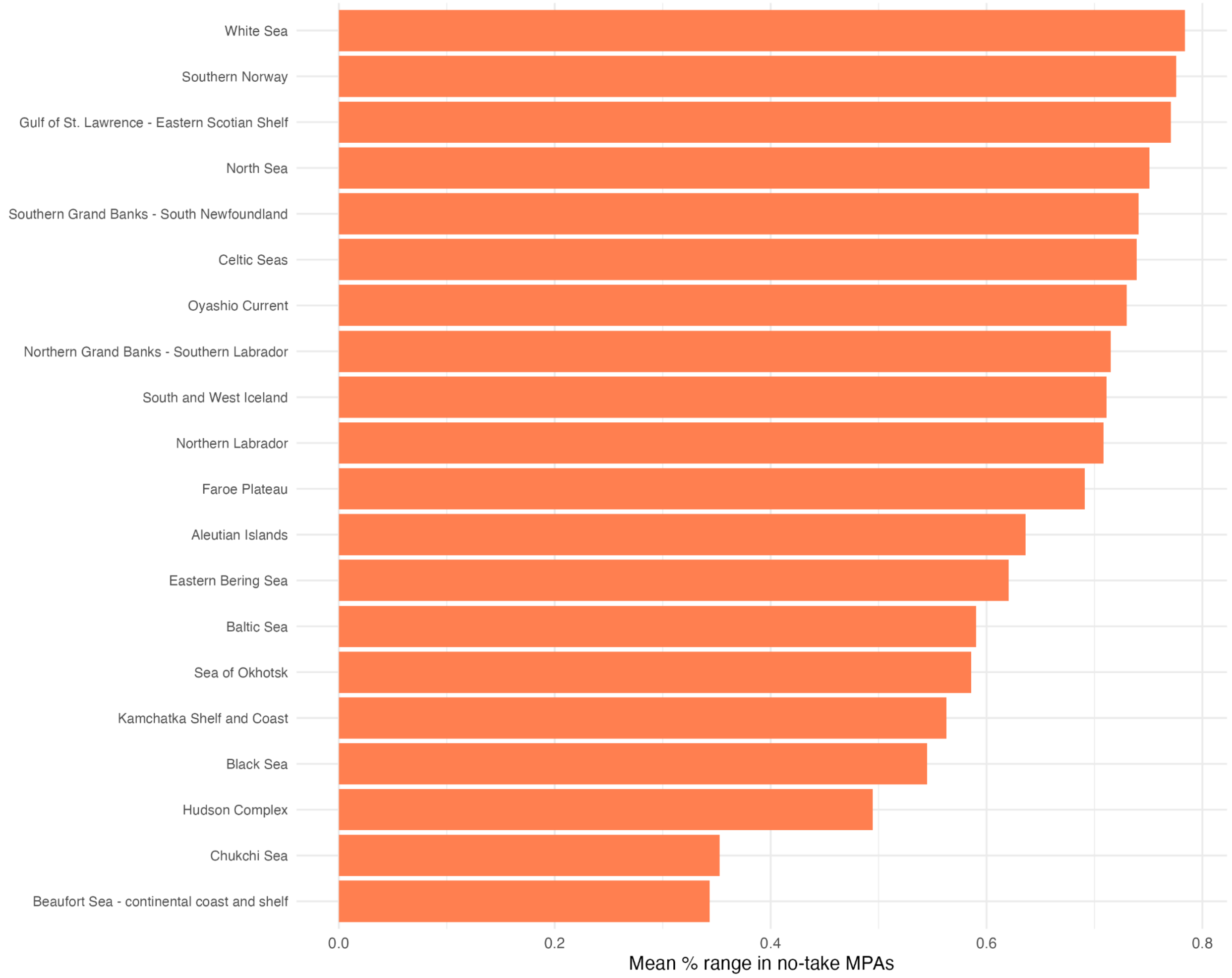


**Fig S4. Bottom 20 marine ecoregions ranked by mean percentage range overlapped by no-take marine protected areas.** Bar chart showing the marine ecoregions with the lowest mean percentage of species ranges protected within no-take marine protected areas (MPAs). Ecoregions are ranked in ascending order of mean coverage. These ecoregions represent areas with the most limited no-take MPA protection relative to marine biodiversity, predominantly concentrated in Arctic waters and enclosed or semi-enclosed seas.

**Table S2. Top 10 elasmobranch species with the greatest over- and under-representation in no-take Marine Protected Areas relative to null model expectations.** Species are ranked by their Standardized Effect Size (SES), with positive values indicating over-representation (species have more range overlap with no-take MPAs than expected by chance) and negative values indicating under-representation (species have less overlap than expected). "Actual %" shows the observed percentage of each species' range within no-take MPAs, "Random %" shows the mean expected percentage based on 100 null model iterations, and "Difference" shows the absolute percentage point difference between observed and expected values. Only species with SES values beyond ±1.96 ($p < 0.05$) are included. The total number of significantly over- and under-represented species is provided in the note below the table.

Top 10 significantly over/under-represented species in no-take MPAs

| Species | Actual % | Random % | Difference | SES |
|---|---|---|---|---|
| **Over-represented** | | | | |
| Narke capensis | 0.78 | 0.00 | 0.78 | Inf |
| Pseudoginglymostoma brevicaudatum | 0.98 | 0.00 | 0.98 | Inf |
| Trigonognathus kabeyai | 44.27 | 2.60 | 41.67 | 32.80 |
| Apristurus spongiceps | 28.74 | 2.44 | 26.30 | 26.41 |
| Parmaturus albimarginatus | 5.00 | 0.03 | 4.97 | 19.90 |
| Parmaturus albipenis | 5.56 | 0.06 | 5.50 | 14.07 |
| Etmopterus villosus | 34.21 | 2.29 | 31.92 | 13.91 |
| Mobula tarapacana | 1.15 | 0.97 | 0.18 | 10.35 |
| Hemiscyllium galei | 36.36 | 1.36 | 35.00 | 9.97 |
| Carcharhinus galapagensis | 6.30 | 2.97 | 3.33 | 9.56 |
| **Under-represented** | | | | |
| Carcharhinus obscurus | 3.13 | 6.05 | -2.92 | -13.82 |
| Hypogaleus hyugaensis | 5.69 | 16.07 | -10.38 | -13.69 |
| Sphyrna lewini | 1.58 | 2.99 | -1.41 | -13.66 |
| Carcharias taurus | 2.04 | 6.09 | -4.05 | -13.49 |
| Sphyrna mokarran | 1.41 | 2.58 | -1.17 | -12.15 |
| Carcharhinus plumbeus | 1.82 | 3.32 | -1.50 | -11.26 |
| Sphyrna zygaena | 0.99 | 2.68 | -1.69 | -11.18 |
| Etmopterus lucifer | 1.92 | 5.25 | -3.33 | -11.18 |
| Amblyraja hyperborea | 1.01 | 2.37 | -1.35 | -10.75 |
| Heterodontus portusjacksoni | 4.04 | 21.28 | -17.25 | -10.72 |

*Note:*

Total significantly over-represented species: 133

Total significantly under-represented species: 365

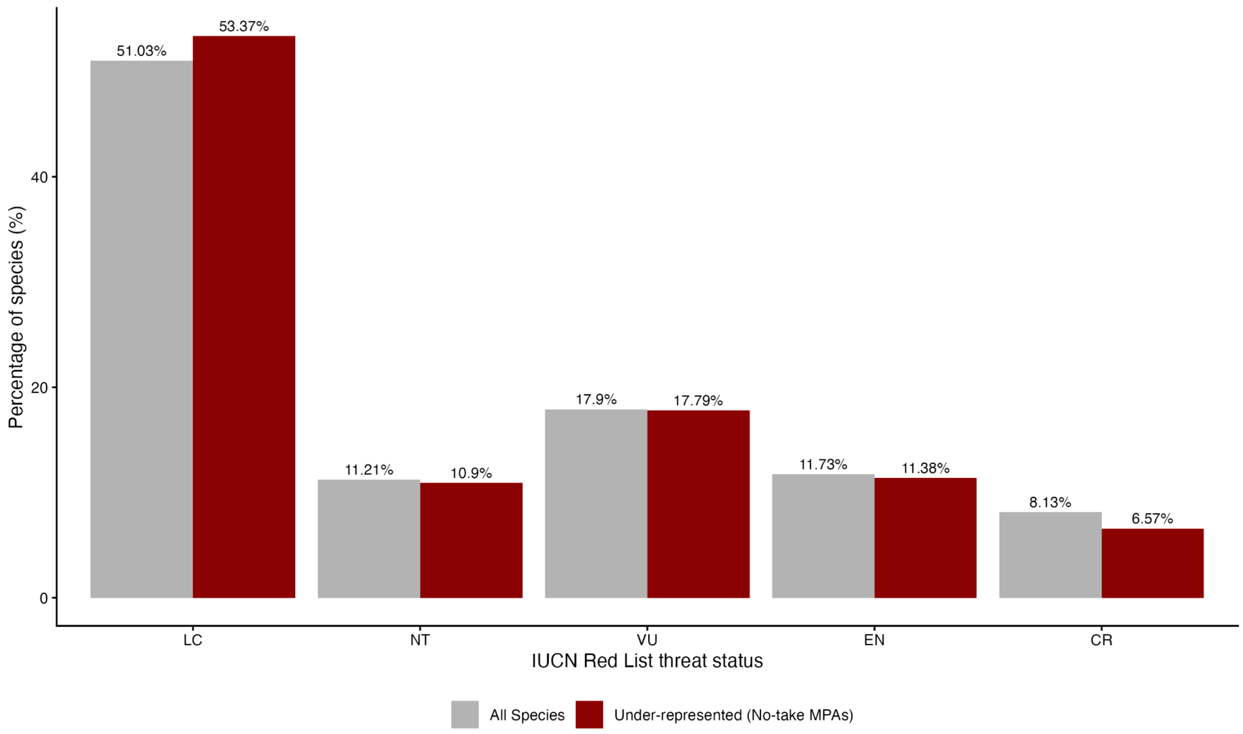


**Fig S5. IUCN threat status distribution of elasmobranch species under-represented in no-take MPAs compared to all studied species.** Bar chart comparing the percentage of species in each IUCN Red List threat category (LC=Least Concern, NT=Near Threatened, VU=Vulnerable, EN=Endangered, CR=Critically Endangered) between all species analysed (grey bars) and species under-represented in no-take MPAs (dark red bars). Under-represented species are those with actual no-take MPA coverage less than would be expected from random MPA placement. Values above each bar indicate the percentage of species within each threat category for the respective group.

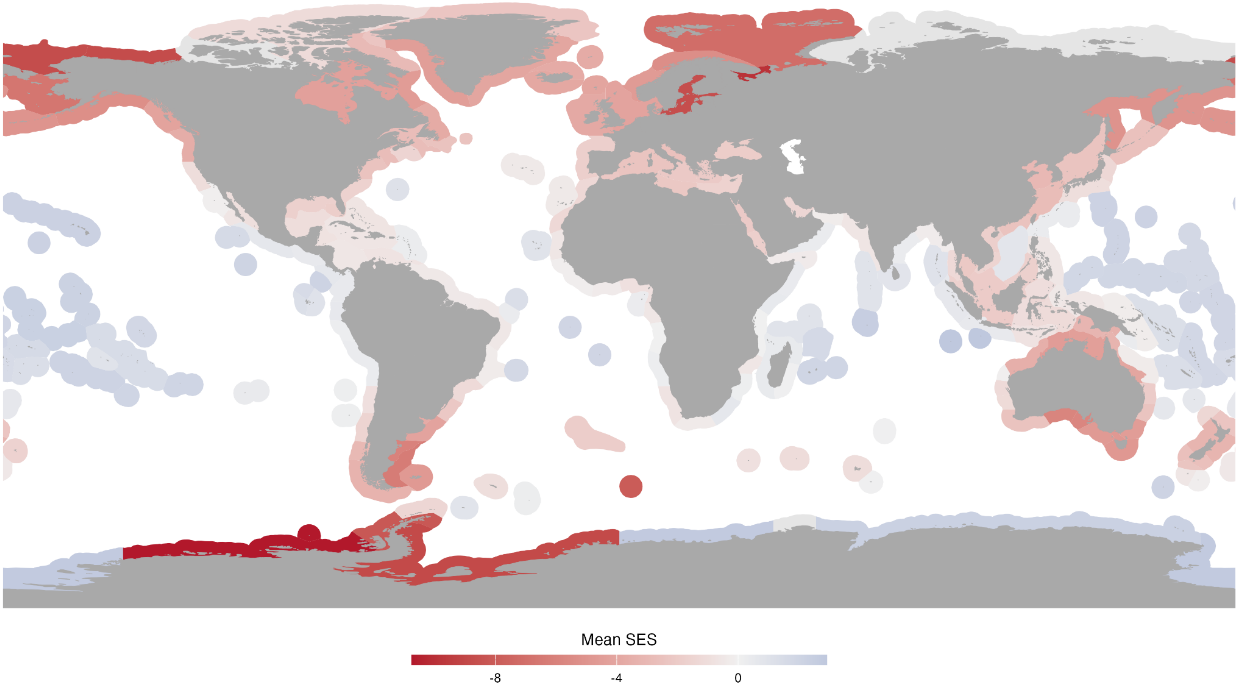


**Fig S6. Standardized effect size (SES) of observed versus null no-take marine protected area coverage by marine ecoregion.** The map displays mean SES values across the world's marine ecoregions, based on the Marine Ecoregions of the World (MEOW) classification system. SES values indicate whether observed no-take MPA coverage within each ecoregion is significantly higher (blue, positive values) or lower (red, negative values) than expected under a null model of random MPA placement. Values near zero (white) indicate that observed coverage does not differ substantially from random expectations. Grey areas represent ecoregions with insufficient data. Land areas are shown in dark grey for reference. The analysis reveals substantial biogeographic variation in the effectiveness of current no-take MPA placement relative to biodiversity conservation priorities.

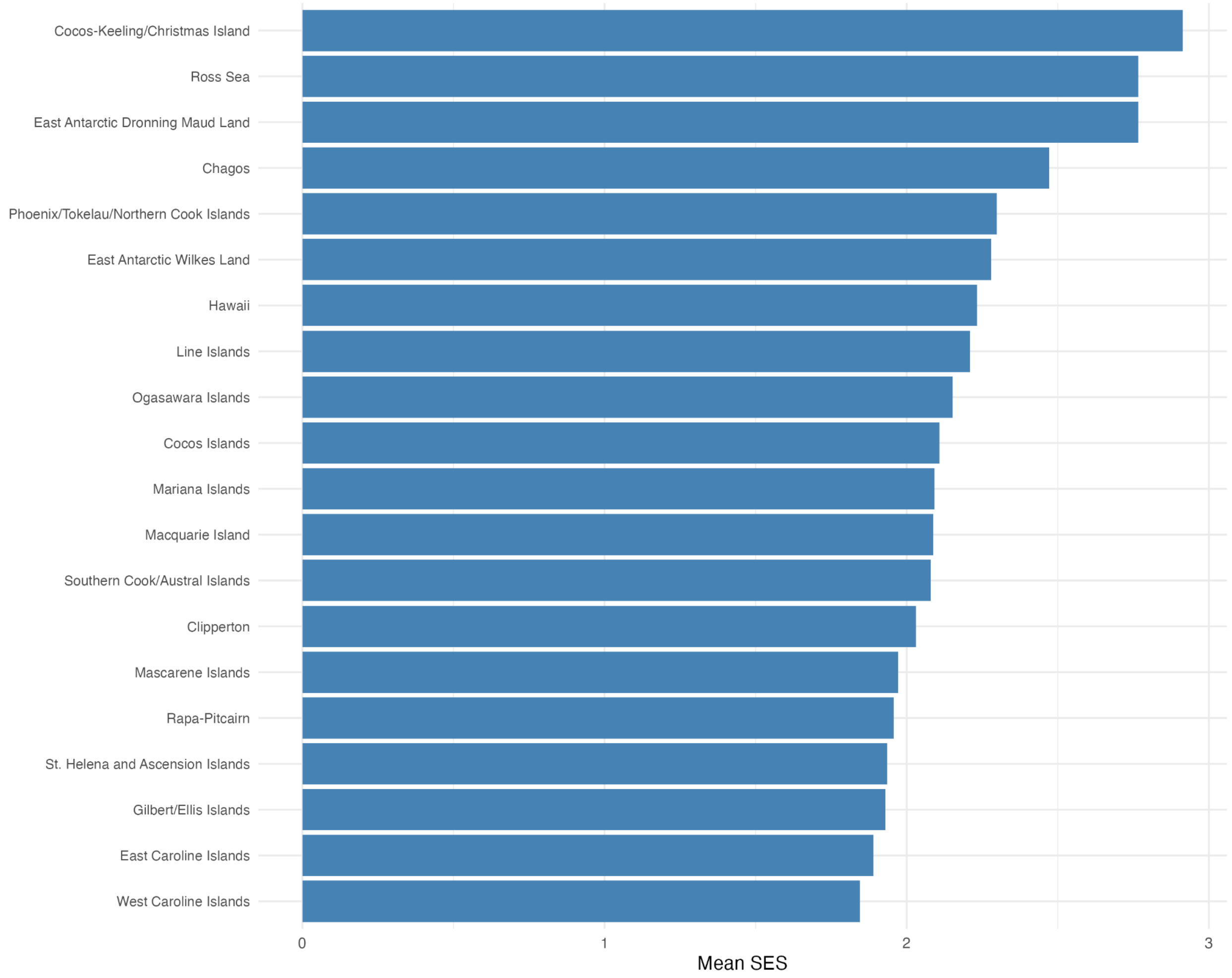


**Fig S7. Top 20 marine ecoregions with highest mean standardized effect sizes (SES) for no-take marine protected area coverage.** Bar chart showing the marine ecoregions with the most positive SES values, indicating that observed no-take MPA coverage significantly exceeds what would be expected under random placement. Ecoregions are ranked in ascending order of mean SES. These ecoregions demonstrate the most effective targeting of no-take MPAs relative to biodiversity conservation priorities, with observed coverage substantially higher than null model expectations.

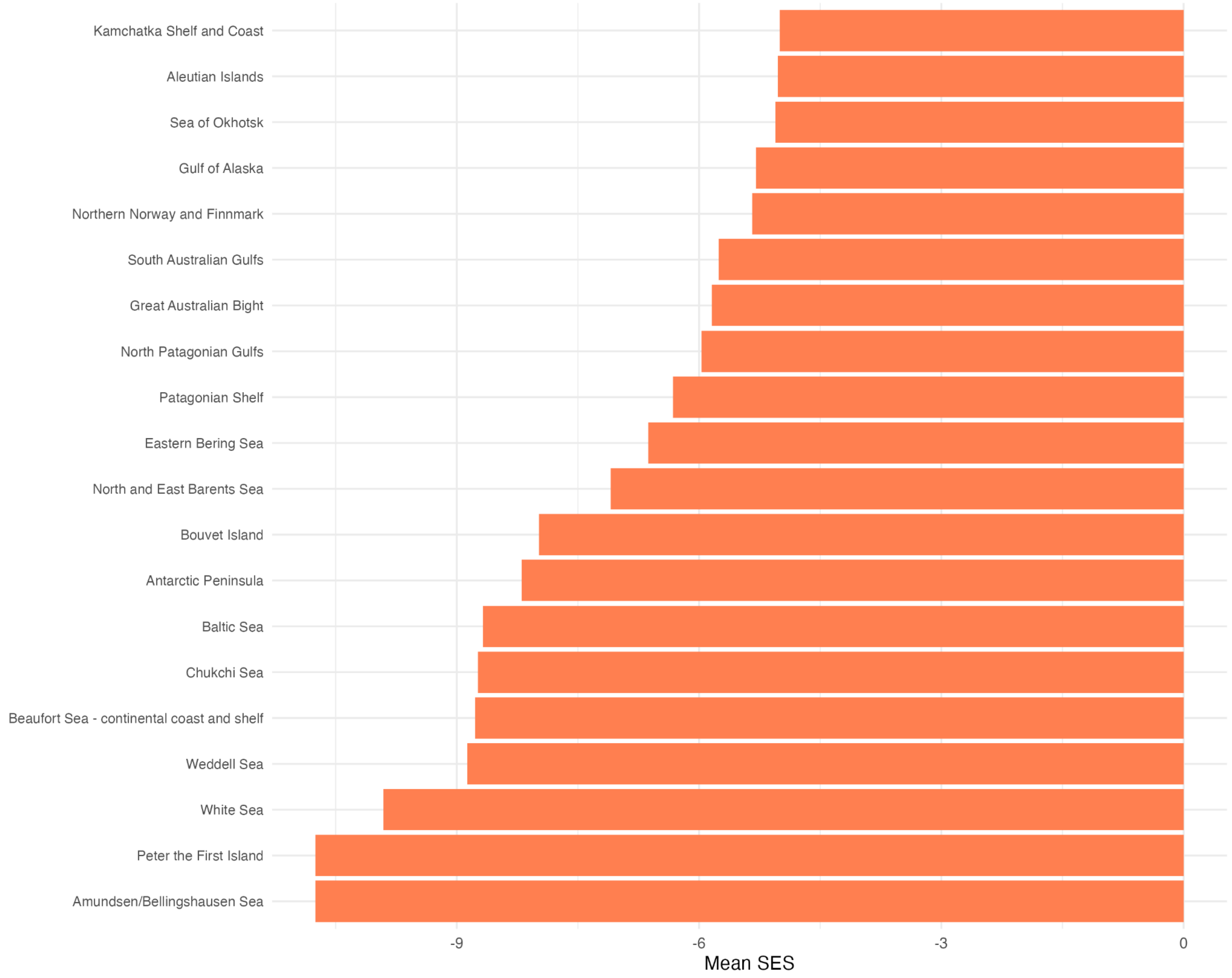


**Fig S8. Bottom 20 marine ecoregions with lowest mean standardized effect sizes (SES) for no-take marine protected area coverage.** Bar chart showing the marine ecoregions with the most negative SES values, indicating that observed no-take MPA coverage is significantly lower than what would be expected under random placement. Ecoregions are ranked in ascending order of mean SES.

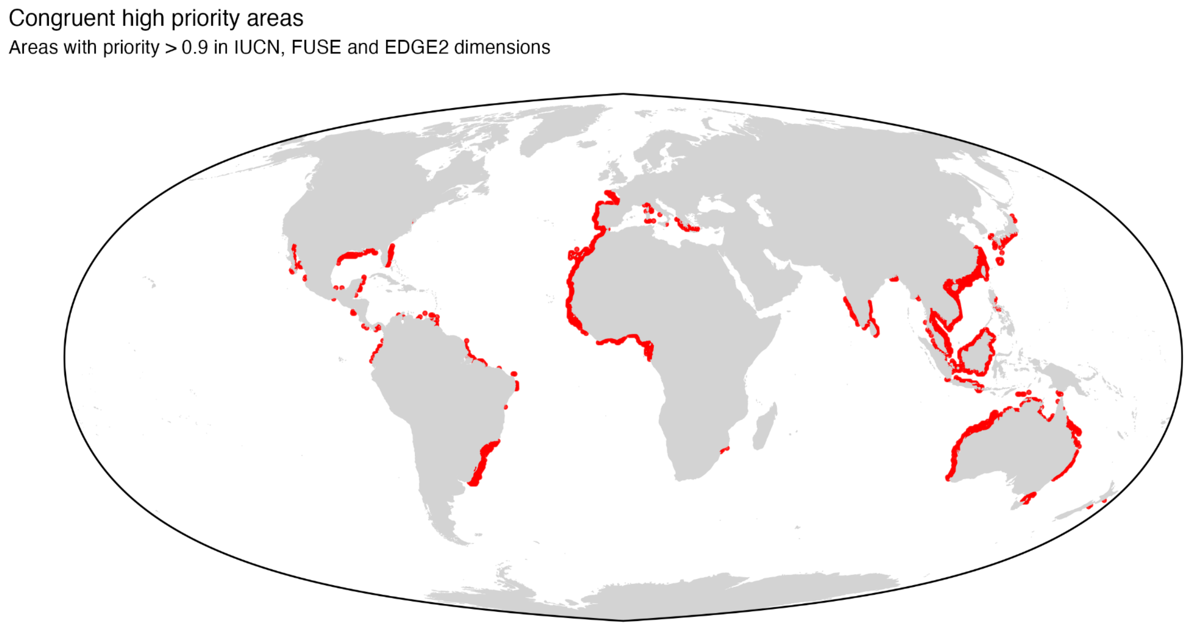


**Fig S9. Spatial congruence of high-priority conservation areas across three dimensions of conservation values.** Red areas indicate planning units where all three dimensions (IUCN, FUSE and EDGE2) show priority scores >0.9, representing areas of highest conservation consensus. The map displays only congruent high-priority areas overlaid on a global projection, highlighting regions where IUCN, FUSE and EDGE2-based conservation priorities spatially align. These areas represent conservation opportunities with multiple dimensions of conservation value and may warrant prioritised protection efforts.

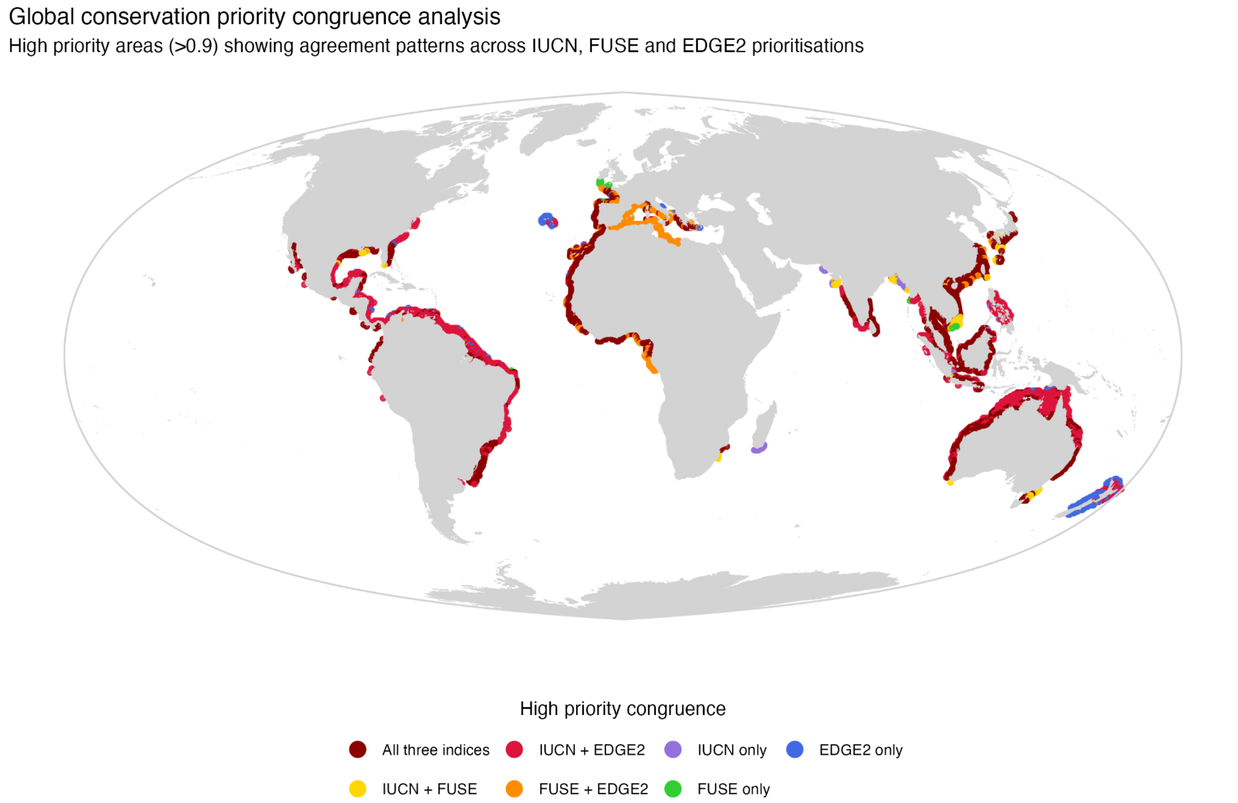


**Fig S10. Global spatial congruence of elasmobranch conservation priorities across three dimensions of conservation value.** The map shows high-priority areas (priority score >0.9) identified by systematic conservation planning using IUCN Red List threat status (IUCN), FUSE and EDGE2 dimensions. Colours indicate agreement patterns: dark red areas represent full congruence across all three dimensions (46.50% of high-priority areas), while other colours show pairwise agreements or dimension-specific priorities. Areas of high congruence primarily occur in coastal waters around Australia, Southeast Asia, Atlantic Africa, European Atlantic coasts, the Caribbean, and South American Atlantic regions. The analysis used a 10% conservation budget constraint with the CAPTAIN algorithm and demonstrates substantial spatial consensus despite methodological differences between prioritisation approaches.

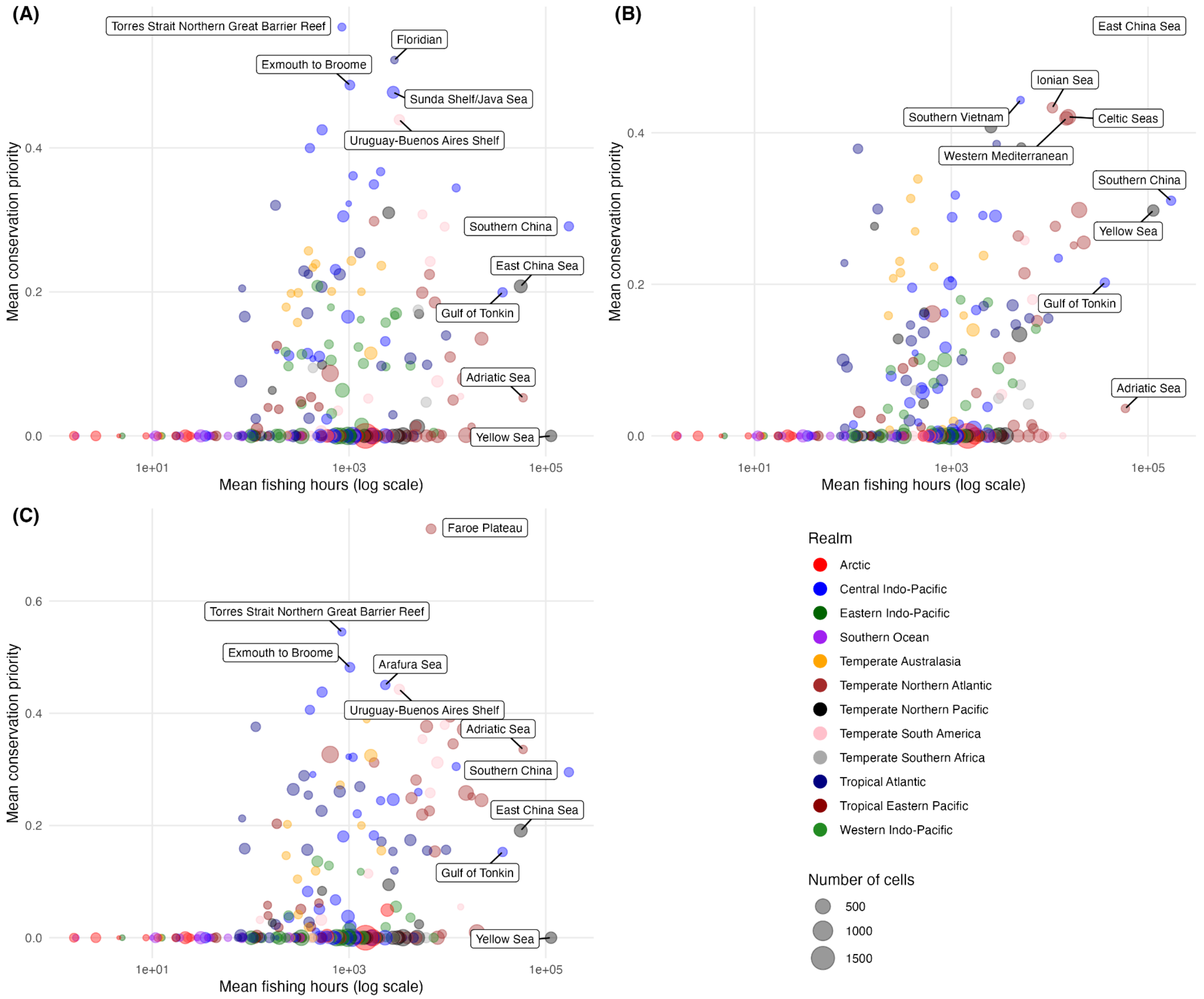


**Fig S11. Relationship between fishing pressure and conservation priority by marine ecoregion.** Scatterplots showing mean fishing hours (log scale) versus mean conservation priority for **(A)** IUCN status, **(B)** FUSE, and **(C)** EDGE2 indices across marine ecoregions. Points are coloured by biogeographic realm and sized by the number of grid cells (data coverage) within each ecoregion. Labels indicate the top 5 ecoregions by fishing pressure and top 5 ecoregions by conservation priority for each dimension. Fishing pressure ranges from 10 to $10^5$ hours, with values $>10^3$ hours representing high fishing intensity. Ecoregions in the upper right quadrants represent conservation conflicts (high priority, high fishing pressure), while those in the upper left represent conservation opportunities (high priority, low fishing pressure).

### Supplementary methods

#### SM1. Predicting fishing activity using random forest modelling

To estimate fishing effort in areas where only Synthetic Aperture Radar (SAR) data was available, we developed a random forest regression model using a comprehensive dataset of over 160,000 matched observations where both Automatic Identification System (AIS) and SAR data were present. The model was implemented at a 0.1-degree spatial resolution global grid. We selected predictor variables that influence fishing distribution patterns: SAR vessel presence scores, geographical coordinates (latitude and longitude), distance to nearest port, distance to shore, and ocean bathymetry. The bathymetric and distance raster layers were resampled to match our 0.1-degree analysis grid.

We evaluated four model variations with different data transformations: (1) no transformations, (2) log-transformed fishing hours, (3) log-transformed SAR presence scores, and (4) log-transformation of both variables. For each model, we used 500 decision trees and assessed variable importance. Model performance was evaluated using multiple metrics including mean absolute error (MAE), root mean squared error (RMSE), mean absolute percentage error (MAPE), median absolute error, R-squared, and adjusted R-squared values. The training and validation datasets consisted exclusively of grid cells where both AIS and SAR detections were present, ensuring direct comparability between observed and predicted fishing effort values.

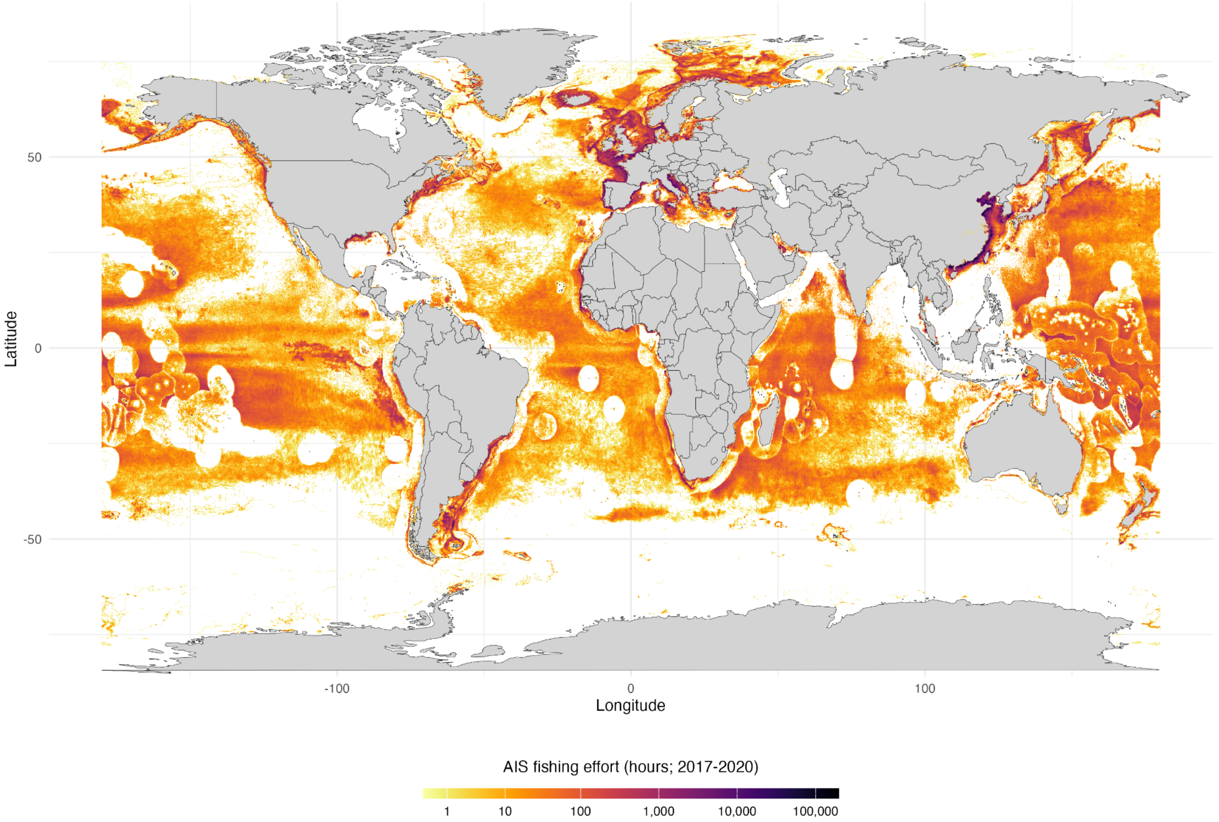


**Fig SM1. Global distribution of industrial fishing effort detected by Automatic Identification System (AIS).** This map shows the spatial distribution of cumulative fishing activity hours from 2017 to 2020, based on vessel tracking data from the Global Fishing Watch database (Kroodsma et al., 2018). Fishing effort is aggregated at 0.1-degree grid resolution and displayed using a logarithmic colour scale (inferno palette), with darker colours representing higher fishing intensity (in hours). The data depicts vessels equipped with AIS transponders, which primarily captures large-scale industrial fishing operations. This dataset provides a comprehensive global view of the footprint of trackable industrial fishing activities, though it may under-represent smaller vessels and those operating without active AIS transmissions. Land masses are shown in light grey.

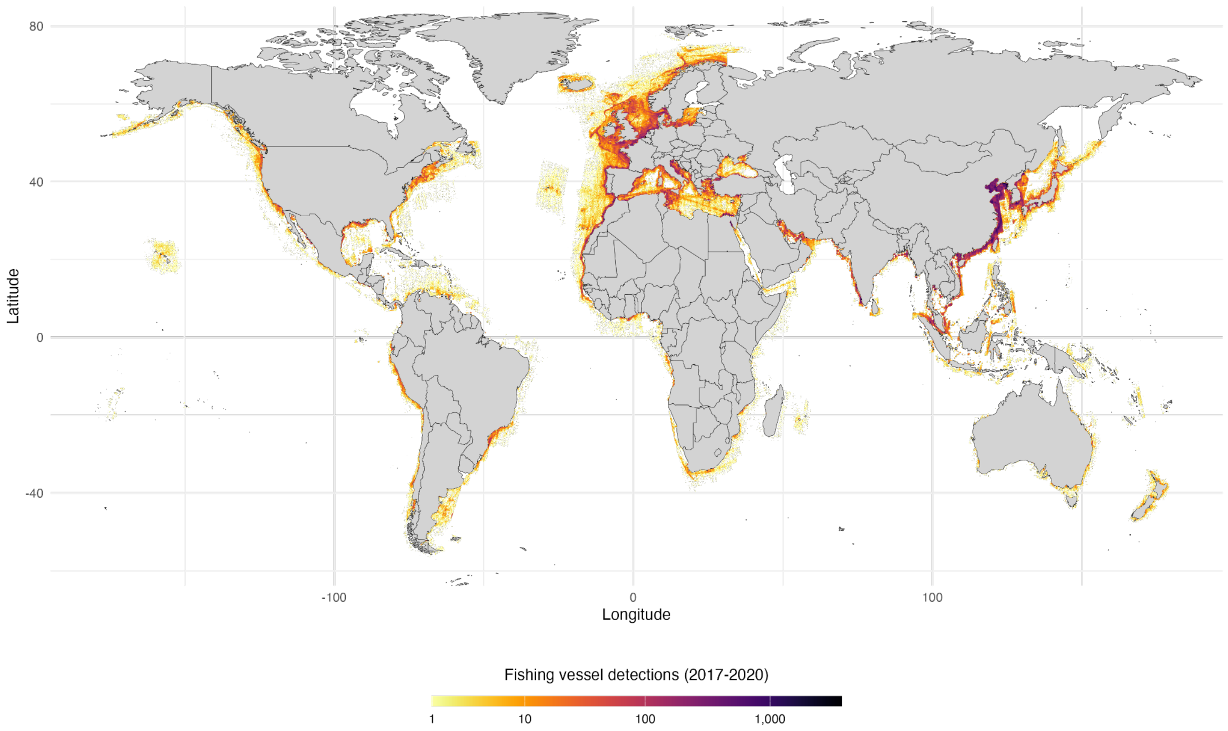


**Fig SM2. Global distribution of fishing vessel detections using Sentinel-1 Synthetic Aperture Radar (SAR).** This map visualizes the spatial distribution of vessel detections with high fishing probability (fishing score ≥ 0.9) during 2017-2020, based on data from the Global Fishing Watch SAR vessel detection program (Paolo et al., 2024). The data is aggregated at 0.1-degree grid resolution, with colour intensity (inferno palette) representing the logarithm of cumulative vessel presence scores. SAR technology can detect vessels regardless of whether they broadcast AIS signals, thereby capturing both compliant and potentially non-compliant fishing activities. This dataset complements AIS-based observations by detecting vessels that might otherwise be invisible to tracking systems, providing a more comprehensive picture of global fishing activity, particularly in regions with higher rates of unmonitored or unreported fishing. Land masses are shown in light grey.

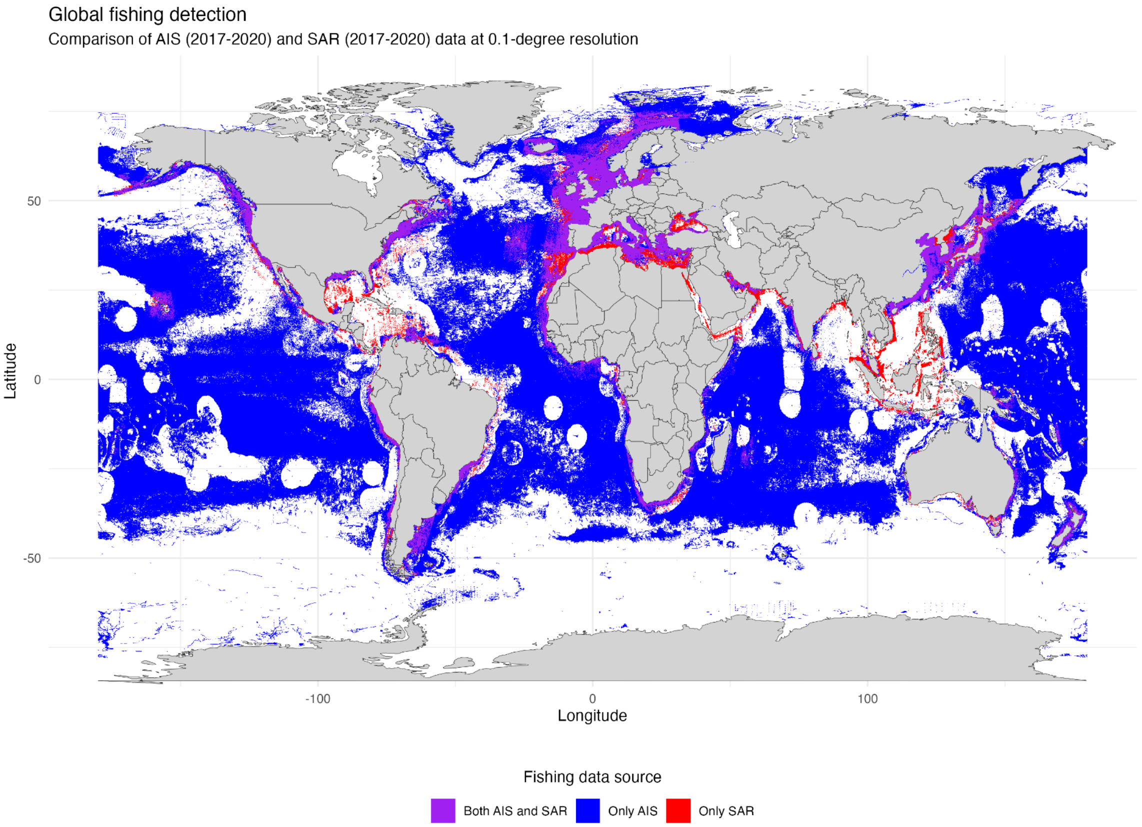


**Fig SM3. Spatial comparison of fishing activity detected by AIS and SAR monitoring systems (2017-2020).** This map illustrates the global distribution of fishing activities as captured by two complementary detection methods: Automatic Identification System (AIS) and Synthetic Aperture Radar (SAR). Areas where fishing was detected by both technologies are shown in purple. Blue areas indicate locations where only AIS signals were detected. Red areas highlight locations where vessels were detected only by SAR. Data is aggregated at 0.1-degree grid resolution over a four-year period (2017-2020). This comparative visualization demonstrates substantial differences in spatial coverage between the two monitoring systems and underscores the importance of using multiple detection technologies to comprehensively map global fishing activities. The analysis reveals significant fishing activities that would remain invisible if relying solely on AIS data, particularly in certain coastal waters. Land masses are shown in light grey.

**Table SM2. Distribution of fishing activity detection across monitoring technologies.** This table quantifies the relative coverage of fishing activity detections by Automatic Identification System (AIS) and Synthetic Aperture Radar (SAR) technologies from 2017 to 2020. The "Category" column identifies three distinct detection scenarios: areas where both technologies detected fishing activity, areas where only AIS signals were recorded, and areas where only SAR detected vessels. The "Number of cells" column presents the count of 0.1-degree grid cells falling into each category, while the "Percentage (%)" column shows the proportion of the total monitored ocean area corresponding to each category. This quantitative breakdown reveals the complementary nature of these monitoring systems and highlights significant spatial differences in coverage, with particular emphasis on the proportion of fishing activity detected exclusively through SAR that would otherwise remain unobserved using AIS monitoring alone.

Summary statistics of data categories

| Category | Number of cells | Percentage (%) |
|---|---|---|
| Both AIS and SAR | 163095 | 9.12 |
| Only AIS | 1566190 | 87.60 |
| Only SAR | 58668 | 3.28 |

#### SM2. Random forest model performance

Our random forest model successfully established quantitative relationships between SAR vessel detections and fishing effort across global marine areas. Among the four model variants tested, the log-transformed fishing hours model demonstrated superior performance with the highest explanatory power ($R^2 = 0.824$, Adjusted $R^2 = 0.824$) and relatively balanced error metrics across different fishing intensity levels. This model maintained significantly lower mean absolute percentage error (69.71%) compared to untransformed models (>1200%), indicating substantially better performance when predicting across the wide range of fishing intensity values present in global fisheries.

The model identified SAR vessel presence scores, geographical location, and bathymetry as the strongest predictors of fishing intensity, which aligns with established understanding of fishing distribution patterns. Distance to port and shoreline also contributed meaningful predictive power. The logarithmic transformation of fishing hours effectively addressed the right-skewed distribution of fishing effort data, resulting in more balanced prediction accuracy across both high and low fishing intensity areas. This model configuration was subsequently applied to predict fishing hours in global ocean grid cells where only SAR detections were available, substantially expanding the spatial coverage of quantified fishing effort estimates beyond what AIS data alone could provide.

**Table SM3. Performance comparison of random forest models with different data transformations for predicting fishing effort.** This table presents eight performance metrics for four random forest model variations tested for predicting fishing hours from SAR vessel detections and environmental variables. Each column represents a different data transformation approach: no transformations ("No transform"), log-transformed fishing hours only ("Fishing hours log"), log-transformed SAR presence scores only ("Presence score log"), and log-transformation of both variables ("Both log"). The metrics include error measurements (Mean absolute error, Root mean squared error, Mean absolute percentage error, Median absolute error), model fit statistics (R-squared, Adjusted r-squared), and residual characteristics (mean and standard deviation). The fishing hours log model demonstrates superior overall performance with the highest r-squared value (0.824) and substantially lower mean absolute percentage error (69.7%) compared to untransformed models, indicating better predictive accuracy across the wide range of global fishing intensity values. This model was selected for the final prediction of fishing hours in areas with SAR-only detection.

Model performance comparison

| | Models | | | |
|---|---|---|---|---|
| **Metric** | **No transform** | **Fishing hours log** | **Presence score log** | **Both log** |
| **Mean absolute error** | 160.27 | 186.46 | 160.91 | 186.25 |
| **Root mean squared error** | 737.54 | 1123.12 | 742.74 | 1119.79 |
| **Mean absolute percentage error** | 1228.12 | 71.13 | 1242.68 | 71.20 |
| **Median absolute error** | 21.97 | 10.10 | 22.01 | 10.10 |
| **R-squared** | 0.81 | 0.82 | 0.81 | 0.82 |
| **Adjusted R-squared** | 0.81 | 0.82 | 0.81 | 0.82 |
| **Mean of residuals** | -5.94 | 150.59 | -6.44 | 150.29 |
| **Standard deviation of residuals** | 737.52 | 1112.98 | 742.72 | 1109.66 |

**Table SM4. Performance metrics of the selected log-transformed fishing hours random forest model.** This table presents detailed evaluation metrics for the final selected random forest model that uses log-transformed fishing hours as the target variable while using untransformed SAR presence scores and environmental predictors. The metrics quantify the model's prediction accuracy and explanatory power when validated against known AIS fishing hours data. The model demonstrates strong explanatory power with an R-squared value of 0.82, indicating it explains approximately 82% of the variance in fishing effort across global marine areas. Error metrics show a mean absolute error of 92.58 hours and median absolute error of 10.19 hours, with a mean absolute percentage error of 69.71%. The near-zero mean of residuals (0.01) indicates minimal systematic bias in predictions. This model was selected for implementation to estimate fishing hours in areas where only SAR vessel detections were available, enabling comprehensive global fishing effort mapping by integrating both monitoring technologies.

Model evaluation metrics for log-transformed fishing hours model

| Metric | Value |
|---|---|
| Mean absolute error | 186.46 |
| Root mean squared error | 1123.12 |
| Mean absolute percentage error | 71.13 |
| Median absolute error | 10.10 |
| R-squared | 0.82 |
| Adjusted r-squared | 0.82 |
| Mean of residuals | 150.59 |
| Standard deviation of residuals | 1112.98 |

**Table SM5. Variable importance metrics for the log-transformed fishing hours random forest model.** This table ranks predictor variables based on their contribution to model performance in estimating fishing effort from SAR detections. Two complementary importance metrics are shown: %IncMSE (percent increase in mean squared error) indicates the performance decrease when a variable is randomly permuted, while IncNodePurity measures the total decrease in node impurity from splits on each variable across all trees. Variables are sorted in descending order of importance based on %IncMSE.

Feature importance for log-transformed fishing hours model

| | Feature | %IncMSE | IncNodePurity |
|---|---|---|---|
| **lat_std** | lat_std | 298.39 | 23760.26 |
| **lon_std** | lon_std | 243.32 | 26855.30 |
| **bathy** | bathy | 214.46 | 28714.10 |
| **total_presence_score** | total_presence_score | 189.20 | 44184.39 |
| **dist_shore** | dist_shore | 136.41 | 12774.27 |
| **dist_ports** | dist_ports | 99.18 | 14170.55 |

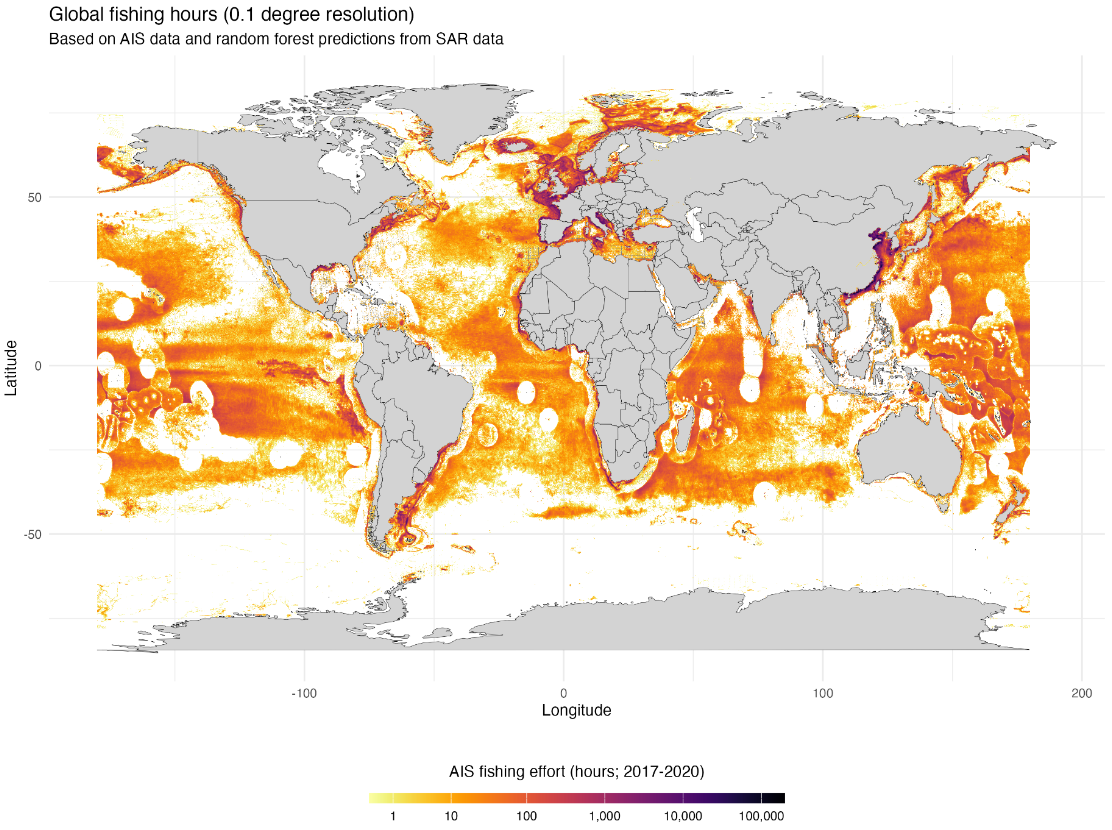


**Fig SM4. Integrated global fishing effort map combining AIS data and random forest model predictions from SAR data (2017-2020).** This map visualizes fishing effort at 0.1-degree resolution by combining direct Automatic Identification System (AIS) observations with random forest model predictions for areas where only Synthetic Aperture Radar (SAR) detections were available. The colour gradient represents fishing hours on a logarithmic scale, spanning from zero (bright yellow) to over 100,000 hours (dark purple) accumulated over the four-year period. The integrated approach reveals substantial fishing activities in regions where AIS coverage alone would indicate minimal or no fishing.
